# A photo-switchable assay system for dendrite degeneration and repair in *Drosophila melanogaster*

**DOI:** 10.1101/2021.08.09.455574

**Authors:** Han-Hsuan Liu, Chien-Hsiang Hsu, Lily Y. Jan, Yuh-Nung Jan

## Abstract

Neurodegeneration arising from aging, injury or disease has devastating health consequences. Whereas neuronal survival and axon degeneration have been studied extensively, much less is known about how neurodegeneration impacts dendrites. To develop an assay for dendrite degeneration and repair in the *Drosophila* peripheral nervous system, we used photo-switchable caspase-3 (caspase-LOV) to induce neuronal damage with tunable severity by adjusting illumination duration, thereby revealing cell type-specific responses to caspase-3 induced dendrite degeneration in dendrite arborization (da) neurons. To ask whether mechanisms underlying axon degeneration also govern dendrite degeneration, we tested the involvement of the Wallerian degeneration pathway by examining the effects of expressing the mouse Wallerian degeneration slow (Wld^S^) protein and knockdown of the *Drosophila* sterile alpha/Armadillo/Toll-Interleukin receptor homology domain protein (dSarm1) and Axundead (Axed) in class 4 da neurons. Here we report Wld^S^ expression or knockdown of dSarm1 improved dendrite repair following caspase- 3 induced dendrite degeneration. Whereas both dSarm1 and Axed were required for thermal nocifensive behavior in uninjured animals, Wld^S^ expression improved the recovery of thermal nocifensive behavior that was impaired by chronic low-level of caspase-LOV activity. By establishing ways to induce graded dendrite degeneration, we uncover a protective role of Wld^S^ in caspase-3 induced dendrite degeneration and repair.

## INTRODUCTION

Neurodegeneration may cause disabilities that place tremendous burdens on both patients and society at large. While much progress has been made in the study of neuronal survival and axon degeneration, it remains an open question as to how dendrites respond to injuries or neurodegeneration. Dendrite degeneration may result from neurological disorders, traumatic brain injury, aging, and other insults (Kulkarni and Firestein, 2012; Kweon et al., 2017; Mulherkar et al., 2017; Penzes et al., 2011; Xiong et al., 2019). These deleterious changes in dendrite structures impair how neurons receive and process information, likely causing major deficits to neurological function (Mulherkar et al., 2017; Penzes et al., 2011). Elucidating the underlying mechanisms of dendrite degeneration and repair will help to uncover ways to reduce damage and facilitate recovery and thus has important clinical implications. Physiologically relevant and reliable *in vivo* injury models are key to better understanding how dendrite degeneration may be reduced and to what extent dendrites are capable of repair.

*Drosophila* dendrite arborization (da) neurons are well suited for studying dendrite development, degeneration, and repair. They are sensory neurons in the body wall and the confinement of their dendrites in a primarily two-dimensional space is conducive to live imaging (Jan and Jan, 2010). Based on the dendrite arbor complexity, da neurons are grouped into four classes with class 4 da (c4da) neurons displaying the most complex dendrite arbors (Grueber et al., 2002). Da neurons can sense and initiate response to different harmful sensory modalities. For example, c4da neurons can detect high temperature, harsh mechanical stimulation, noxious chemicals, and harmful short wave-length light (Gorczyca et al., 2014; Hwang et al., 2012; Kim et al., 2012; Xiang et al., 2010; Zhong et al., 2010), whereas class 3 da (c3da) neurons are specialized for sensing gentle mechanical stimulation (Yan et al., 2013). The behavioral readouts of da neurons, such as the fast crawling and rolling escape behaviors initiated by c4da neurons upon high temperature, are well-characterized and can be used for assessments of functional recovery (Babcock et al., 2009; Hwang et al., 2007). Studies that use laser ablation to sever dendrites from the c4da, c3da and c1da neuron somata have shown that dendrites can repair themselves. The repair process depends on kinases, electrical activity, extracellular environment, microRNA, and kinetochore proteins (DeVault et al., 2018; Hertzler et al., 2020; Kitatani et al., 2020; Nye et al., 2020; Song et al., 2012; Stone et al., 2014; Thompson-Peer et al., 2016). However, the harsh injury caused by dendrite severing is likely more severe and drastic as compared to insults induced by neurological disorders, traumatic brain injury, aging, and other insults. Moreover, laser ablation is labor-intensive and hence not suitable for high-throughput screening designed to uncover novel mechanisms. In order to gain insights on how dendrites degenerate and repair, it is desirable to develop an alternative neurodegeneration model that can better simulate how a neuron responds to the insults that it may encounter in its lifetime.

Many conditions can induce neurodegeneration. In this study, we used caspase-3, which acts downstream of various insults, as a switch to initiate neurodegeneration. Activation of caspase-3, an executor for apoptotic cell death, has been observed in neurons exposed to insults such as injury, neurotoxins, and neurodegenerative diseases (Cotman and Su, 1996; Eldadah and Faden, 2000). There are also circumstances where, following caspase-3 activation, neurons stay alive and display degeneration or partial remodeling in dendrites or axons (Erturk et al., 2014; Khatri et al., 2018; Kuo et al., 2006; Simon et al., 2016; Williams et al., 2006). These observations suggest that caspase-3 could be used as a way to introduce damage on dendrites systematically to elicit neurodegeneration. A recently developed photo-switchable caspase-3, caspase-LOV, provides opportunities to test whether a controllable caspase-3 could be a versatile tool to induce neurodegeneration with diverse outcomes ranging from apoptosis to repair (Smart et al., 2017). In this system, a light-oxygen-voltage-sensing domain (LOV domain) is inserted into the intersubunit linker of human caspase-3 (Smart et al., 2017). Illumination with 450 nm light activates this photo- switchable caspase-3, and the activation only lasts for the duration of illumination. This reversible feature of caspase-LOV makes it possible to adjust the degree of caspase-3 activity during a specific time window (Smart et al., 2017).

Wallerian degeneration is an evolutionarily conserved process to clear distal axons after axon injury. This process can be delayed by neuronal expression of the mouse Wallerian degeneration slow (Wld^S^) protein in both mice and flies (Hoopfer et al., 2006; Lunn et al., 1989; MacDonald et al., 2006). Wld^S^ can also partially protect axon degeneration following trophic deprivation and dendrite pruning during metamorphosis, both of which are caspase-3 dependent (Schoenmann et al., 2010; Tao and Rolls, 2011). Caspase-3 independent dendrite degeneration induced by injury or phosphatidylserine (PS) exposure could be delayed with Wld^S^ as well (Ji et al., 2021; Sapar et al., 2018). Interestingly, a study using both mouse and *Drosophila* models raises the possibility that Wld^S^ and caspase act in parallel during dendrite pruning, because Wld^S^ does not supress caspase activity (Schoenmann et al., 2010). Loss-of-function mutations in *Drosophila* Toll receptor adaptor proteins, the sterile alpha/Armadillo/Toll-Interleukin receptor homology domain protein (dSarm1) and Axundead (Axed), both of which are involved in the Wallerian degeneration pathway, afford protection for axon degeneration induced by injury but not axon degeneration during developmental pruning or apoptotic cell death (Neukomm et al., 2017; Osterloh et al., 2012). The suppression of injury-induced axon degeneration can be achieved by knocking down expression of dSarm1 with RNAi as well (Gerdts et al., 2013). Deletion of dSarm1 protects injury- and PS-induced dendrite degeneration (Ji et al., 2021), whereas deletion of Axed only partially protects the injury-induced dendrite degeneration but does not affect PS-induced dendrite degeneration (Ji et al., 2021). It is unclear whether these proteins involved in the Wallerian degeneration pathway play any roles in caspase-3 dependent dendrite degeneration and repair.

To elucidate the cellular mechanism of dendrite degeneration and repair, we used the photo-switchable caspase-3 to induce varying degrees of dendrite degeneration in *Drosophila* larval da neurons and monitored the repair process afterward. We found that the caspase-3 dependent dendrite degeneration in da neurons was worsened by prolonging the illumination, and dendrite repair was evident with attenuated activation of caspase-3. We observed cell type-specific responses to caspase-3 induced dendrite degeneration in da neurons. Expression of mouse Wld^S^ in c4da neurons resulted in longer and more numerous dendrites during caspase-3 induced dendrite degeneration and during development as well. Similarly, knockdown of dSarm1 or Axed, two factors involved in Wallerian degeneration, increased survival of neurons following caspase-LOV activation. Additionally, knockdown of dSarm1 led to longer dendrites both during development and following caspase-LOV activation. Reduced expression of Axed did not affect the dendrite structure during development or following caspase-LOV activation. We further showed that the compromised thermal nocifensive behavior caused by chronic low-level of caspase-LOV activity in c4da neurons can be partially rescued with Wld^S^ expression but not with knockdown of dSarm1 or Axed.

## RESULTS

### Caspase-LOV activation of different durations initiates graded dendrite degeneration in sensory neurons

Among larval da neurons, c4da neurons display the most complex dendrite structures (Grueber et al., 2002). Their dendrites actively grow in length, scale in size to extend coverage area, and continue adding new tips throughout larval development (Grueber et al., 2002; Parrish et al., 2009; Williams and Truman, 2005). In this study, we sought to determine to what extent c4da neurons can recover from caspase-3 induced degeneration following transient activation of a photo-switchable caspase-3, caspase-LOV. The amount of illumination is known to correlate with the amount of caspase-3 activity which can effectively induce dendrite degeneration followed by apoptosis in several type of cells including c4da neurons (Smart et al., 2017).

Given that the activation of caspase-LOV can be easily controlled by adjusting the intensity and the duration of illumination, we began our study by monitoring dendrite degeneration following caspase-LOV activation for 2 hours (h), 30 minutes (min), or 10 min. We used a membrane tethered tdTomato (UAS-CD4-tdTOM) driven by the ppk-GAL4 to label the plasma membrane of c4da neurons for visualization of individual dendrite arbors. Freely moving larvae were illuminated for various durations at 48 h after egg laying (AEL) on transparent agar plates and then transferred back to a dark environment. We performed time-lapse imaging to monitor the dendrite structure of the same c4da neuron, ddaC, 24 h and 72 h following illumination with blue LED (***Figure 1A***). The 24 h and 72 h imaging timepoints provide snap shots for the early and late stages of caspase-3 induced dendrite degeneration and subsequent repair, as indications for the acute and continuing response to the degeneration, respectively.

**Figure 1.**
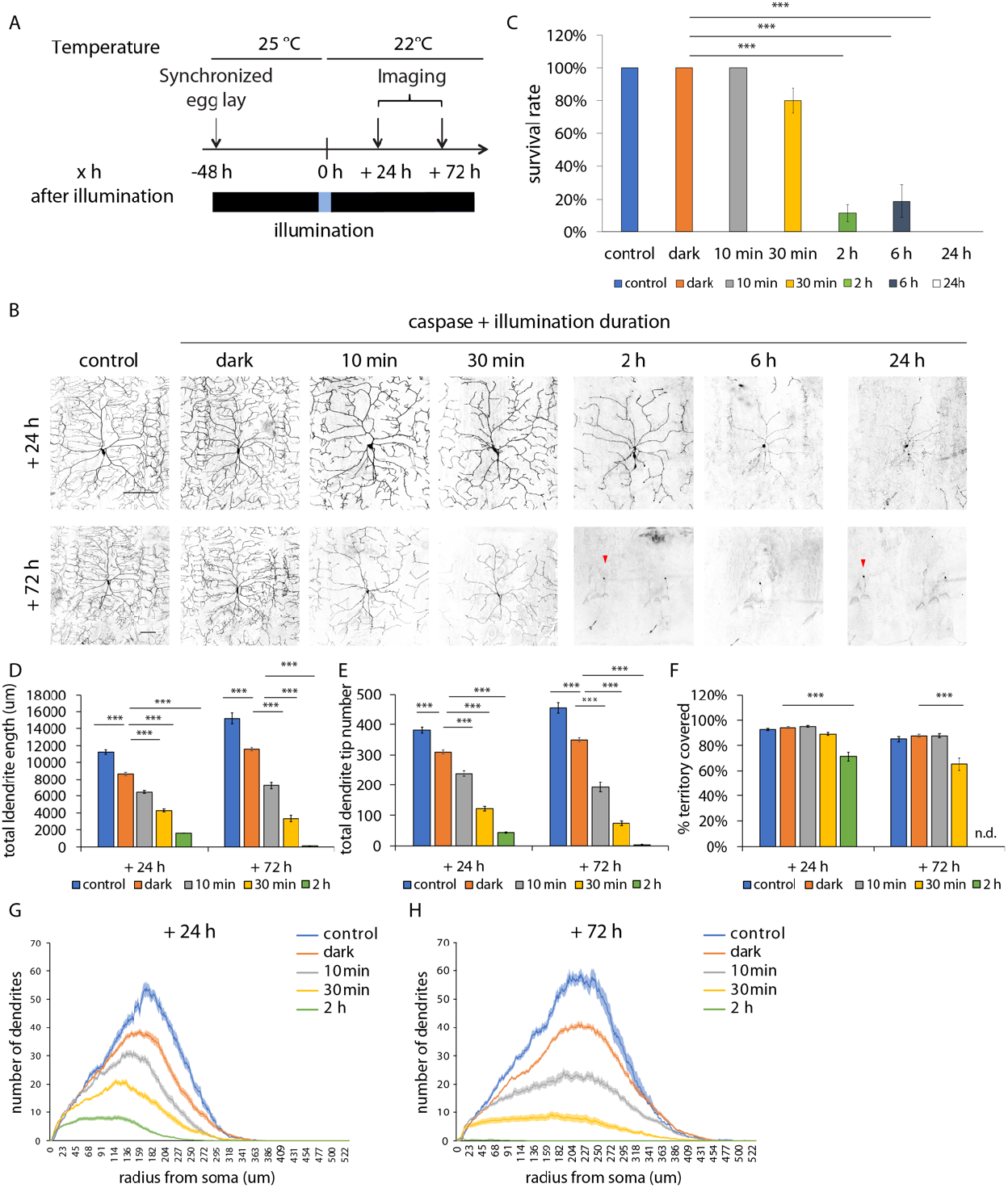
Transient caspase-LOV activation initiates dendrite degeneration followed by repair in c4da neurons. (A) Protocol to illuminate and image larval c4da neurons expressing just UAS-tdTOM (control) or UAS-tdTOM and UAS-caspase-LOV (dark, 10 min-24 h) using ppk-GAL4. These neurons were labeled with tdTOM for visualization. Larvae were kept in the dark all the time (control, dark) or kept in the dark and illuminated at 48 h after egg laying for 10 min-24 h. The same neurons were imaged twice at 24 h and 72 h following illumination. (B) Representative images of c4da neurons from larva without caspase-LOV and kept in the dark (control), with caspase-LOV and kept in the dark (dark), or with caspase-LOV and illuminated for different durations (10 min-24 h). Neurons were imaged at 24 h (+ 24 h, top row) and 72 h (+ 72 h, bottom row) after illumination started. (C) Survival rates of c4da neurons expressing caspase-LOV decrease when illumination is extended. Survival of neurons was counted 72 h after illumination. (D-F) Quantifications of dendrite structures of survived c4da neurons following caspase-LOV activation, including total dendrite length (D), total dendrite tip numbers (E), and percentage of territory covered (F). The skeletal dendrite structures were predicted by in-house built deep learning models. The quantifications were carried out using a python script. (G-H) Sholl analysis of dendrite complexity 24 h (G) and 72 h (H) after illumination. The complexity of the dendrite structure, quantified as numbers of dendrites crossing continuous circles originated from the soma and represented by the total area under the curve, decreases with caspase-LOV expression and progresses as illumination extends. All conditions are significantly different from each other (p<0.01). Scale bars =100 μm. * p<0.05, ** p<0.01, *** p<0.001, Kruskal-Wallis rank sum test with Dunn’s post hoc test further adjusted by the Benjamini-Hochberg FDR method for multiple independent samples (C); one-way ANOVA with Tukey’s post hoc test for multiple comparisons in (D-H). Error bars represent ± SEM (C-F) or in shaded area (G-H). n = 14-55 neurons for each experimental condition and timepoint.

To facilitate the quantification of complex morphology of c4da neurons in this study, we built a deep learning model based on the U-Net architecture (Ronneberger et al., 2015) which has been widely used for biomedical image segmentation, including detecting dendrite branch terminals of da neurons (Kanaoka et al., 2019). We applied our model to automatically segment dendrite structure from microscopy images and retrieve segmentation masks containing the full reconstruction of the dendrite arbors of neurons. Segmentation masks of individual neurons were then used to measure different parameters of neuronal morphology, including total dendrite length, total dendrite tip numbers, percentage of territory covered, and dendrite complexities assessed with Sholl analysis. We validated that the dendrite structures segmented by our model and found that they were comparable to manual reconstruction and achieved high Dice coefficient, a commonly used spatial overlap index for evaluating segmentation quality (Zou et al., 2004) (***Figure 1 – figure supplement 1A***). To further evaluate the model performance, we compared parameters of neuronal morphology measured from model-predicted segmentation with those derived from manual reconstruction by using the images of c4da neurons acquired in Figure 1. With post-processing to fill in gaps and remove small fragments (see Methods), we observed high correlation for both tip numbers (R^2^ = 0.97) and total dendrite length (R^2^ = 0.99; ***Figure 1 – figure supplement 1B***,***C***).

Caspase-LOV activation lasting longer than 2 h induced apoptosis within 72 h (***Figure 1B***,***C***). Shortening the caspase-LOV activation to 30 min allowed the average survival rate for illuminated neurons to reach 80%. With 10 min caspase-LOV activation, almost all neurons survived for at least 72 h (***Figure 1C***). Using the deep learning-based model, we quantified the dendrite structures of c4da neurons that were either kept in the dark or illuminated for a duration ranging from 10 min to 2 h. Activation of caspase-LOV for 2 h caused the reduction of total dendrite length, tip numbers, dendrite complexity, and percentage of territory covered, both at 24 h and at 72 h following caspase-LOV activation (***Figure 1B***,***D***,***E***,***F***,***G***,***H***). The total dendrite length, tip numbers, and dendrite complexity decreased progressively with increasing durations of illumination, while the percentage of territory covered was affected at 72 h after 30 min illumination (***Figure 1B***,***D***,***E***,***F***,***G***,***H***). Neurons exposed to 30 min of blue LED illumination displayed significantly shorter and fewer dendrites compared to those exposed to 10 min illumination. The basal activity of caspase-LOV in the dark (dark) led to reduced dendrite arbor length, tip numbers, and dendrite complexity at the 24 h and 72 h timepoints compared to the animals without caspase-LOV expression (control) (***Figure 1B***,***D***,***E***,***G***,***H***). The percentage of territory covered by dendrites is not affected by caspase-LOV expression if the animals were kept in the dark (***Figure 1F***). With 30 min and 2 h of caspase-LOV activation, there were overall reductions in both dendrite length and tip numbers (***Figure 2A***,***B***). Interestingly, there were still increases in the dendrite length 24-72 h after the 10 min illumination (***Figure 2A***), even though the total dendrite tip numbers were reduced (***Figure 2B***), suggesting that c4da neurons can continue to grow after experiencing caspase-LOV activation. These changes could be a combination of normal dendrite growth, dendrite degeneration, and repair.

**Figure 2.**
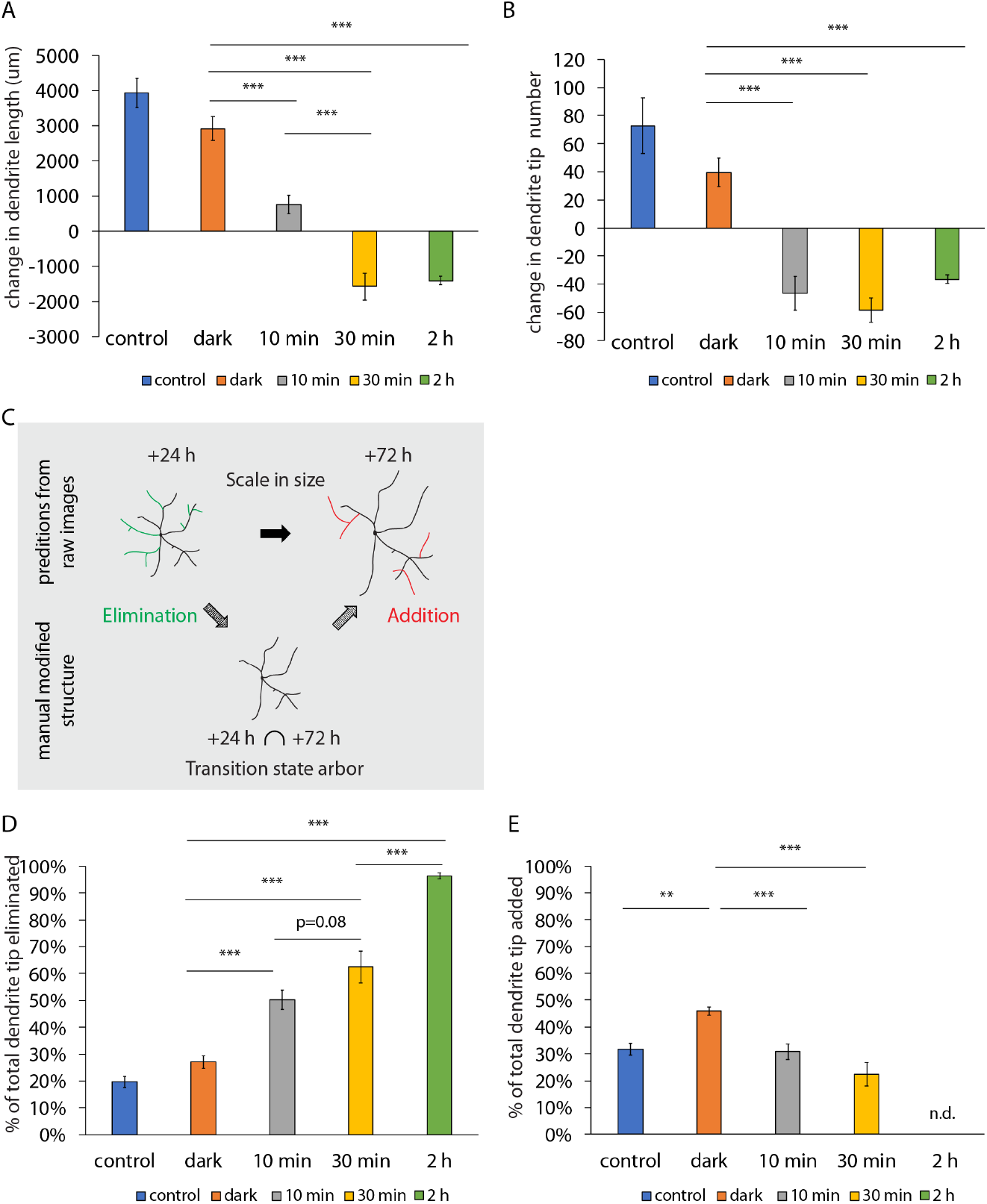
Dendrite addition and elimination occurs simultaneously during the repair process. (A-B) Quantifications of changes in dendrite length (A) and dendrite tip numbers (B) of c4da neurons during the 24 h to 72 h time period after caspase-LOV activation. C4da neurons expressing caspase-LOV decrease growth in dendrite length and dendrite tip numbers as illumination is extended (C) Illustration of elimination and addition of dendrites happened over the degeneration and repair process. (D-E) Quantifications for the percentage of eliminated (D) and added (E) dendrite tips over the 24 h to 72 h time period following caspase-LOV activation. The percentage of tips eliminated increases with longer illumination while the percentage of tips added decreases. * p<0.05, ** p<0.01, *** p<0.001, one-way ANOVA with Tukey’s post hoc test for multiple comparison in (A-B, D-E). Error bars represent ± SEM. n = 19-23 neurons for each experimental condition and timepoint.

To further examine the dendrite elimination and addition of c4da neurons, we analyzed the dendrite dynamics in the tip numbers over a period of 48 h following caspase-LOV activation. We compared the dendrite structure between the 24 h and 72 h timepoints and used the dendrite arbor at 24 h following illumination as the backbone to generate a “transition state arbor” which contained only dendrites observed at both 24 h and 72 h. Then, we subtracted the number of tips of the “transition state arbor” from that at 24 h to give a measure of the eliminated dendrites (those dendrite branches only observed at 24 h), and from that at 72 h to give a measure of the newly added branches (those dendrite branches only observed at 72 h) (***Figure 2C***). The percentage of eliminated dendrite tips was calculated by dividing the number of eliminated dendrites by the total number of dendrite tips measured at 24 h. The percentage of added dendrite tips was calculated by dividing the number of newly added dendrites by the total number of dendrite tips measured at 72 h.

We found that dendrite elimination and addition took place concurrently in individual neurons following caspase-LOV activation (***Figure 2D***,***E***). As the duration of Caspase-3 activity increased, the percentage of eliminated dendrite tips increased and the percentage of added dendrite tips decreased. Interestingly, even though caspase-LOV activation for 30 min caused reduction in total dendrite length and tip numbers, there were still new branches added following dendrite degeneration. The reduction in total tip numbers following 10 min or 30 min illumination (***Figure 2B***) resulted from the significantly greater increase in elimination (***Figure 2D***) than addition of dendrite branches (***Figure 2E***). C4da neurons expressing caspase-LOV but kept in the dark were comparable with c4da neurons not expressing caspase-LOV based on the percentage of eliminated dendrite tips (***Figure 2D***) though the former had a higher percentage of newly added dendrites (***Figure 2E***).

Taken together, we found that neurons can survive 10-30 min of caspase-LOV activation through illumination, and their dendrites continue to grow in length with addition of new tips to the remaining dendrite arbors. Most of the neurons failed to survive following caspase-LOV activation for longer than 2 h and showed severe dendrite degeneration before dying. By making use of the varying levels of degeneration induced by different durations of illumination, we can search for machineries used for neuroprotection to improve dendrite degeneration, repair or neuronal survival following caspase-3 induced degeneration.

### Class I ddaE neurons can withstand transient caspase-LOV activation and repair dendrite damage

Class I da (c1da) neurons and class 3 da (c3da) neurons differ from c4da neurons in dendritic morphology, growth dynamics and physiological function. To ask whether their response to caspase-3 induced regeneration and repair is also different from that of c4da neurons, we first examined c1da neurons, which have the simplest dendrite arbor among all classes of da neurons. C1da neurons establish their dendrite arbor early in development and only extend existing branches in length without adding new branches in late larval development (Grueber et al., 2002; Williams and Truman, 2005). They are able to initiate regeneration after dendrotomy as are c4da neurons (Sugimura et al., 2003; Tao and Rolls, 2011; Thompson-Peer et al., 2016).

To assess how c1da neurons would react to caspase-3 induced degeneration, we labeled the c1da ddaE neuron with UAS-CD4-tdTOM driven by the GAL4^2-21^ (Grueber et al., 2003a) and used the same paradigm described in ***Figure 1A***. C1da neurons can survived 30 min activation of caspase-LOV, whereas about 10% of c1da neurons imaged were found dead 72 h following caspase-LOV activation for 2 h (***Figure 3A***,***B***). Caspase-LOV activity in the dark (dark) significantly reduced dendrite length at 72 h after illumination (***Figure 3C***). The dendrite tip numbers of c1da neurons expressing caspase-LOV and maintained in the dark (dark) were fewer than those of c1da neurons without caspase-LOV expression (control) (***Figure 3D***). Both 30 min and 2 h caspase-LOV activation impaired dendrite structures (***Figure 3A***,***C***,***D***). Caspase-LOV activation for 2 h induced more drastic reductions in both total dendrite length and tip numbers at 72 h after illumination, as compared to 30 min of caspase-LOV activation (***Figure 3A***,***C***,***D***). The increase in dendrite length (***Figure 3E***) and total tip numbers (***Figure 3F***) over the 24-72 h period following 2 h of caspase-LOV activation was significantly less than dark and 30 min of caspase-LOV activation.

**Figure 3.**
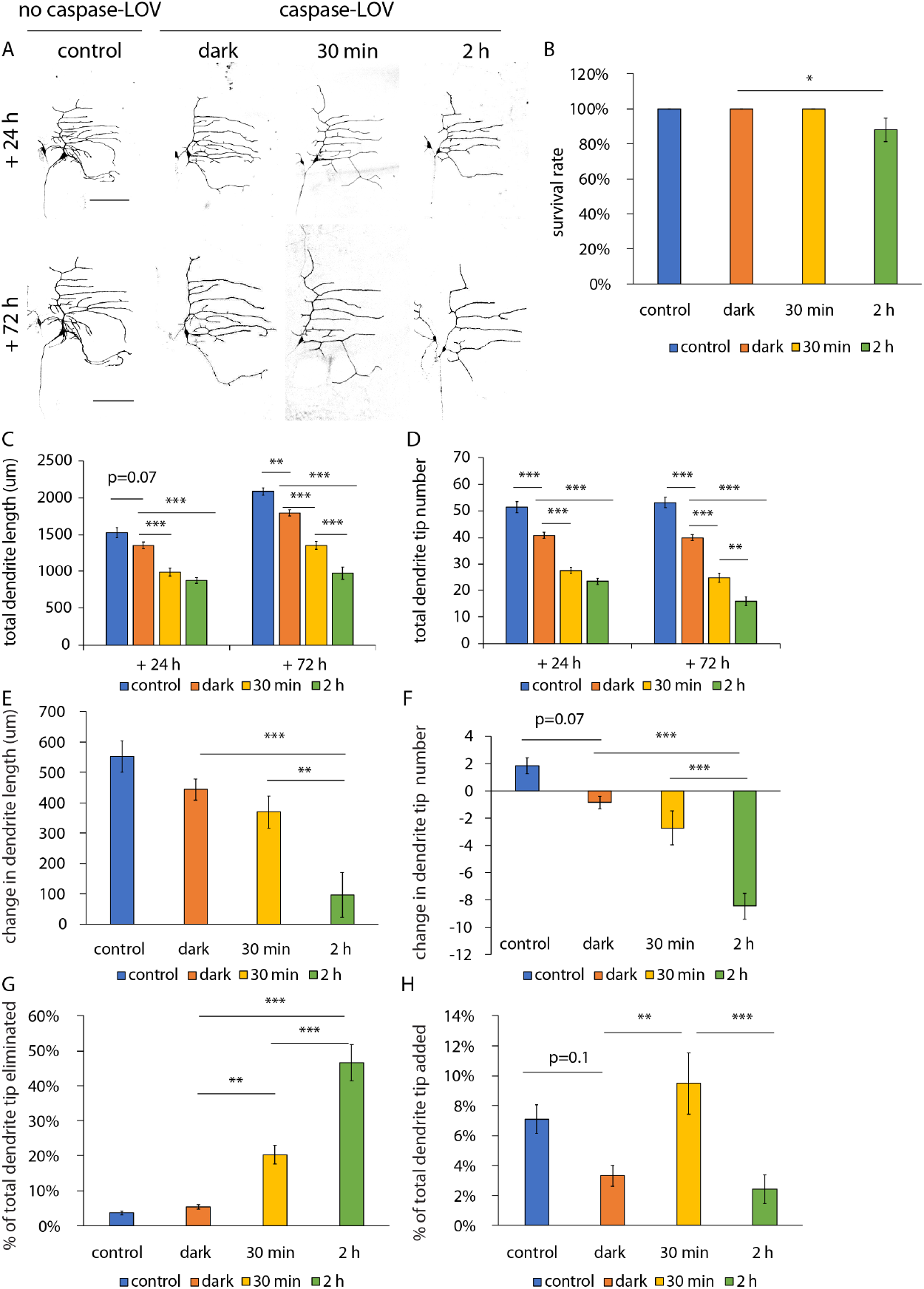
Class I ddaE neurons can sustain mild caspase-LOV activation and repair by adding new branches. (A) Representative images of c1da neurons expressing just UAS-tdTOM (control) or UAS-tdTOM and UAS-caspase-LOV (dark, 30 min, 2 h) driven by ppk^2-21^-GAL4. Larvae were kept in the dark all the time (control, dark) or kept in the dark and illuminated for different durations (30 min, 2 h). The same neurons were imaged at 24 h (top row) and at 72 h (bottom row) after illumination started. (B) Survival rates of c1da neurons are reduced with 2h illumination. About 10% of c1da neuron imaged were found dead 72 h following 2 h caspase-LOV activation. (C-D) Quantifications of dendrite structures of c1da neurons following caspase-LOV activation, including total dendrite length (C) and total dendrite tip numbers (D). (E-F) Quantifications of changes in dendrite length (E) and tip numbers (F) of c1da neurons over the 24 h-72 h time period after caspase-LOV activation. (G-H) Quantifications for the percentage of eliminated (G) and added (H) dendrite tips over the 24 h-72 h time period following caspase-LOV activation. Scale bars =100 μm. * p<0.05, ** p<0.01, *** p<0.001, Kruskal-Wallis rank sum test with Dunn’s post hoc test further adjusted by the Benjamini-Hochberg FDR method for multiple independent samples (B); one-way ANOVA with Tukey’s post hoc test for multiple comparisons in (C-H). Error bars represent ± SEM. n = 22-28 neurons for each experimental condition and timepoint.

We next looked into the dendrite dynamics and quantified dendrite tip elimination and addition as we did for c4da neurons. Similar to previous reports on the limited increase in dendrite tips after early development (Stone et al., 2014; Sugimura et al., 2003), we found that the c1da ddaE neurons without caspase-LOV (control) had 7% and 4% tips added and eliminated, respectively (***Figure 3F***,***G***). Caspase-LOV activity in the dark did not significantly alter the percentage of addition or elimination of dendrite tips. Caspase-LOV activation for 30 min or 2 h increased the percentage of eliminated dendrite tips (***Figure 3G***). There was a robust increase inthe percentage of added dendrite tips of c1da neurons expressing caspase-LOV following 30 min illumination compared to those kept in the dark (***Figure 3H***). This robust increase in the percentage of added dendrite tips was not observed in c1da neurons following 2 h illumination nor in c4dan neurons following any durations of illumination tested in this study (***Figure 2E*** and ***Figure 3H***). Thus, there appears to be a regrowth program unique for c1da neurons that is initiated following 30 min caspase-LOV activation – a program that is not evident following severe degeneration induced by 2 h caspase-LOV activation.

### Class III ddaE neurons can withstand transient caspase-LOV activation induced dendrite damage

C3da neurons have signature bushy tertiary branches enriched in actin (Nagel et al., 2012; Tsubouchi et al., 2012). In contrast to the c4da ddaC and c1da ddaE neurons, c3da neurons do not persist after metamorphosis (Shimono et al., 2009; Williams and Truman, 2005). To test whether they can survive caspase-3 activation, we expressed caspase-LOV in the c3da ddaF neurons and imaged these neurons at 24 and 72 h after illumination. We labeled the c3da ddaF neurons with UAS-CD4-tdTOM driven by the GAL4^19-21^ along with Repo-Gal80 to eliminate the expression in glial cells (Awasaki et al., 2008; Xiang et al., 2010). The c3da ddaF neurons can survive 30 min but not 2 h caspase-LOV activation (***Figure 4A***,***B***). The survival rate of c3da ddaF neurons was significantly reduced to 71% following 2 h caspase-LOV activation (***Figure 4A***,***B***). Caspase-LOV activity in the dark (dark) induced significant reduction in dendrite tip numbers but did not alter the total dendrite length compared to c3da neurons without caspase-LOV (control) (***Figure 4C***,***D***). Both 30 min and 2 h of caspase-LOV activation in c3da ddaF neurons led to reduction in total dendrite length and tip numbers (***Figure 4C***,***D***). Caspase-LOV activity in the dark significantly reduced the increase in dendrite length (***Figure 4E***) but did not significantly alter the dendrite tip numbers (***Figure 4F***). The dendrite length and tip numbers still exhibited increases over the 24-72 h period following 30 min or 2 h of caspase-LOV activation (***Figure 4E***,***F***). Moreover, there were significant increases in the percentage of eliminated tips and significant decreases in the percentage of added new tips following 2 h of caspase-LOV activation (***Figure 4G***,***H***). Thus, c3da ddaF neurons also appear to have a class-specific response to caspase-3 induced dendrite degeneration. They do not initiate regrowth following mild degeneration as observed in c1da ddaE neurons. Instead, there was greater degeneration of c3da neuronal dendrites following longer caspase activation, similar to what we observed in c4da neurons. The c3da ddaF neurons differ from c4da ddaC neurons in that they do not show any increase in the percentage of added dendrite tips with caspase-LOV activity in the dark (***Figure 2E*** and ***Figure 4H***) and they continue to grow in length and add new tips following caspase-LOV activation (***Figure 2A***,***B*** and ***Figure 4E***,***F***).

**Figure 4.**
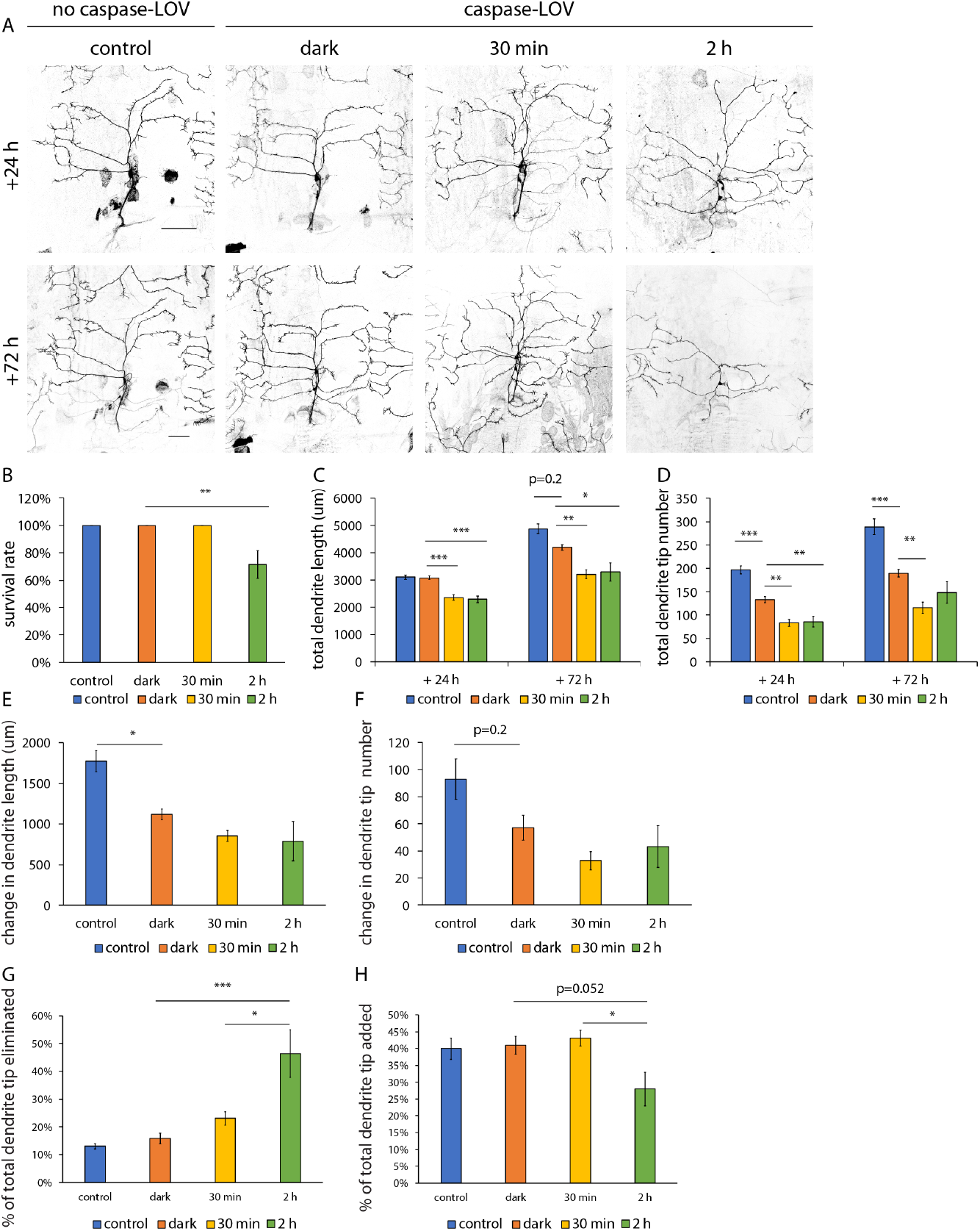
Class III ddaF neurons can sustain mild caspase-LOV activation and repair by adding new branches. (A) Representative images of c3da neurons expressing just UAS-tdTOM (control) or UAS-tdTOM and UAS-caspase-LOV (dark, 30 min, 2 h) driven by ppk^19-12^-GAL4 along with Repo-Gal80. Larvae were kept in the dark all the time (control, dark) or kept in the dark and illuminated for different durations (30min, 2 h). The same neurons were imaged at 24 h (top row) and at 72 h (bottom row) after illumination started. (B) Survival rates of c3da neurons are reduced with 2h illumination. (C-D) Quantifications of dendrite structures of c3da neurons following caspase-LOV activation, including total dendrite length (C) and total dendrite tip numbers (D). (E-F) Quantifications of change in dendrite length (E) and tip numbers (F) of c3da neurons over the 24 h-72 h time period after caspase-LOV activation. (G-H) Quantifications for the percentage of eliminated (G) and added (H) dendrite tips over the 24 h-72 h time period following caspase-LOV activation. Scale bars =100 μm. * p<0.05, ** p<0.01, *** p<0.001, Kruskal-Wallis rank sum test with Dunn’s post hoc test further adjusted by the Benjamini-Hochberg FDR method for multiple independent samples (B); one-way ANOVA with Tukey’s post hoc test for multiple comparisons in (C-H). Error bars represent ± SEM. n = 9-21 neurons for each experimental condition and timepoint.

### Wld^S^ protects c4da neurons from caspase-3 dependent dendrite degeneration

This new degeneration assay system can be used to address questions such as how caspase-LOV activation induces dendrite degeneration, how neurons manage to survive from transient caspase- LOV activation, and how neurons repair their damaged dendrites. We decided to focus on c4da neurons because they have the most complex dendrites among the da neurons (Grueber et al., 2002) and there are established behavioral assays to assess their sensory functions (Babcock et al., 2009; Hwang et al., 2007).

We generated caspase-tester flies expressing the ppk-tdGFP transgene to monitor the dendrite morphology of c4da neurons with caspase-LOV expressed via ppk-GAL4. These tester flies were crossed with either RNAi flies or flies harboring other transgenes of interest. To select illumination conditions, we first examined the degree of dendrite degeneration and repair in c4da neurons labeled with ppk-tdGFP and expressing caspase-LOV and luciferase (control) via ppk-GAL4. We found that 91% of the neurons survived the 10 min illumination, and the survival rate dropped to 22% following 30 min illumination (***Figure 5 – figure supplement 1A***,***B***). We suspected that the lower survival rate following 30 min illumination here compared to Fig. 1 is due to the stronger ppk-Gal4 used for caspase-tester flies, which has an insertion site different from the ppk-Gal4 used in Fig. 1. The 10 min caspase-LOV activation decreased dendrite length and dendrite tip numbers at 24 h and 72 h after illumination and degeneration was worse when activation of caspase-LOV extended to 30 min (***Figure 5 – figure supplement 1A***,***C***,***D***,***E***). The percentage of territory covered was not affected in the neurons that survived the 10 min or 30 min illumination (***Figure 5 – figure supplement 1A***,***E***).

**Figure 5.**
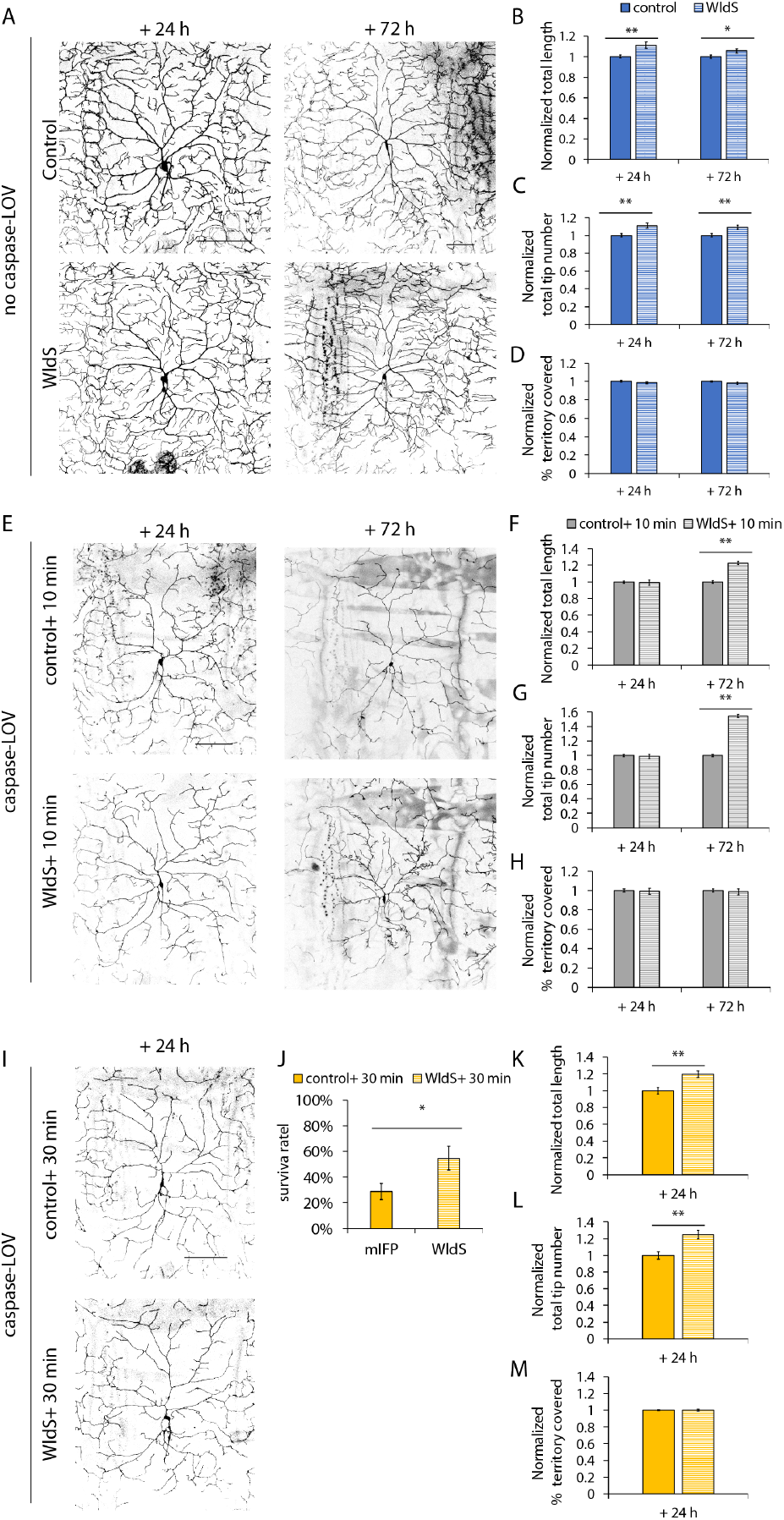
Wld^S^ expressing neurons retain longer and more dendrites during development and upon caspase-3 induced neurodegeneration. (A) Representative images of c4da neurons labeled by ppk-tdGFP with ppk-Gal4 driving expression of UAS-mIFP-2A-HO1 (control) or UAS-Wld^S^ (Wld^S^). (B-D) Quantifications of dendrite structures, including normalized length (B), normalized tip numbers (C), and normalized percentage of territory covered (D) of c4da neurons. (E, I) Representative images of c4da neurons expressing ppk-tdGFP and caspase-LOV with UAS-mIFP-2A-HO1 (control) or UAS-Wld^S^ driven by ppk-Gal4. Larva were kept in the dark and illuminated for 10 min (E) or 30 min (I) at 48 h after egg lay and imaged 24 h or 72 h afterward. (F-H) Quantifications of dendrite structures, including normalized length (F), normalized tip numbers (G), and normalized percentage of territory covered (H) of c4da neurons illuminated for 10 min. (J-M) Quantifications of survival rate (J) and dendrite structures, including normalized length (K), normalized tip numbers (L), and normalized percentage of territory covered (M) of c4da neurons illuminated for 30 min. Scale bars =100 μm. * p<0.05, ** p<0.01, *** p<0.001, Student’s *t* test in (B-D, F-H, K-M), Kruskal-Wallis rank sum test with Dunn’s post hoc test further adjusted by the Benjamini-Hochberg FDR method for multiple independent samples (J); Error bars represent ± SEM. n ≥ 29 neurons for each experimental condition and timepoint.

Wld^S^ has been found to be beneficial for protection against injury-induced dendrite degeneration, PS-induced dendrite degeneration, and developmental dendrite pruning (Ji et al., 2021; Sapar et al., 2018; Schoenmann et al., 2010; Tao and Rolls, 2011). It is unclear whether Wld^S^ is also involved in early dendrite development or caspase-3 dependent dendrite degeneration and repair. During early dendrite development, neurons expressing Wld^S^ displayed mild but significant increases in total dendrite length and tip numbers with no changes in the percentage of territory covered (***Figure 5A***,***B***,***C***,***D***). Using the caspase-tester flies, we examined the functions of Wld^S^ expression in caspase-3 dependent dendrite degeneration and repair. To maintain comparable expression levels of UAS-caspase-LOV driven by ppk-GAL4 in neurons with or without Wld^S^ expression, we include UAS-mIFP-2A-HO1 transgene in the control group. The transgenic flies harboring UAS-mIFP-2A-HO1, which had a wildtype genetic background similar to that of flies with UAS-Wld^S^, expressed monomeric infrared fluorescent proteins (IFP) and Heme Oxygenase 1 Proteins (HO1) driven by Gal4. With caspase-3 induced neurodegeneration, neurons expressing Wld^S^ were comparable to control at 24 h following illumination, but these neurons retained significantly longer dendrites and more numerous dendrite tips at 72 h following 10 min of caspase-LOV activation (***Figure 5E***,***F***,***G***). The protection in dendrite structure afforded by Wld^S^ was already evident at 24 h following 30 min of caspase-LOV activation, as revealed by the longer dendrites and more numerous dendrite tips (***Figure 5I***,***K***,***L***). Wld^S^ expression did not alter the percentage of territory covered following 10 min or 30 min caspase-LOV activation (***Figure 5H***,***M***). With 30 min illumination, neuronal survival was enhanced by Wld^S^ expression in c4da neurons (***Figure 5J***). These results suggest that expression of Wld^S^ can protect c4da neurons from caspase-3 induced dendrite degeneration.

### Knockdown of Axed and dSarm1 are neuroprotective with dSarm1 playing a role in dendrite degeneration and repair

Besides Wld^S^, dSarm1 and Axed are two additional players involved in the Wallerian degeneration pathway. It is unknown how dSarm1 and Axed are involved in early dendrite development and caspase-3 dependent dendrite degeneration and repair in c4da neurons. Hence, we use ppk-GAL4 to drive the expression of luciferase (control) or RNAi targeting dSarm1 or Axed in c4da neurons. During early dendrite development, knocking down dSarm1 in c4da neurons resulted in longer dendrite length without changing the dendrite tip numbers (***Figure 6A***,***B***,***C***,***D***). Knocking down Axed had no significant effect in early dendrite development (***Figure 6A***,***B***,***C***,***D***). During caspase-3 induced dendrite degeneration, neurons with reduced dSarm1 expression had longer and more numerous dendrites and a higher percentage of territory covered with dendrite at 72 h after 10 min illumination (***Figure 6E***,***F***,***G***,***H***). Similar effects on dendrite structure were observed at 24 h following 30 min illumination (***Figure 6I***,***K***,***L***,***M***). Knockdown of Axed did not affect dendrite structure following either 10 min or 30 min illumination (***Figure 6E***,***F***,***G***,***H***,***I***,***K***,***L***,***M***). Neurons with reduced dSarm1 or Axed expression had a higher survival rate (***Figure 6J***). These results indicate that knockdown of dSarm1 or Axed in c4da neurons can increase neuronal survival following caspase-3 induced degeneration, whereas knockdown of dSarm1 but not Axed can protect dendrite structure from caspase-3 induced degeneration. Moreover, dSarm1 is also involved in early dendrite development for regulation of dendrite elongation.

**Figure 6.**
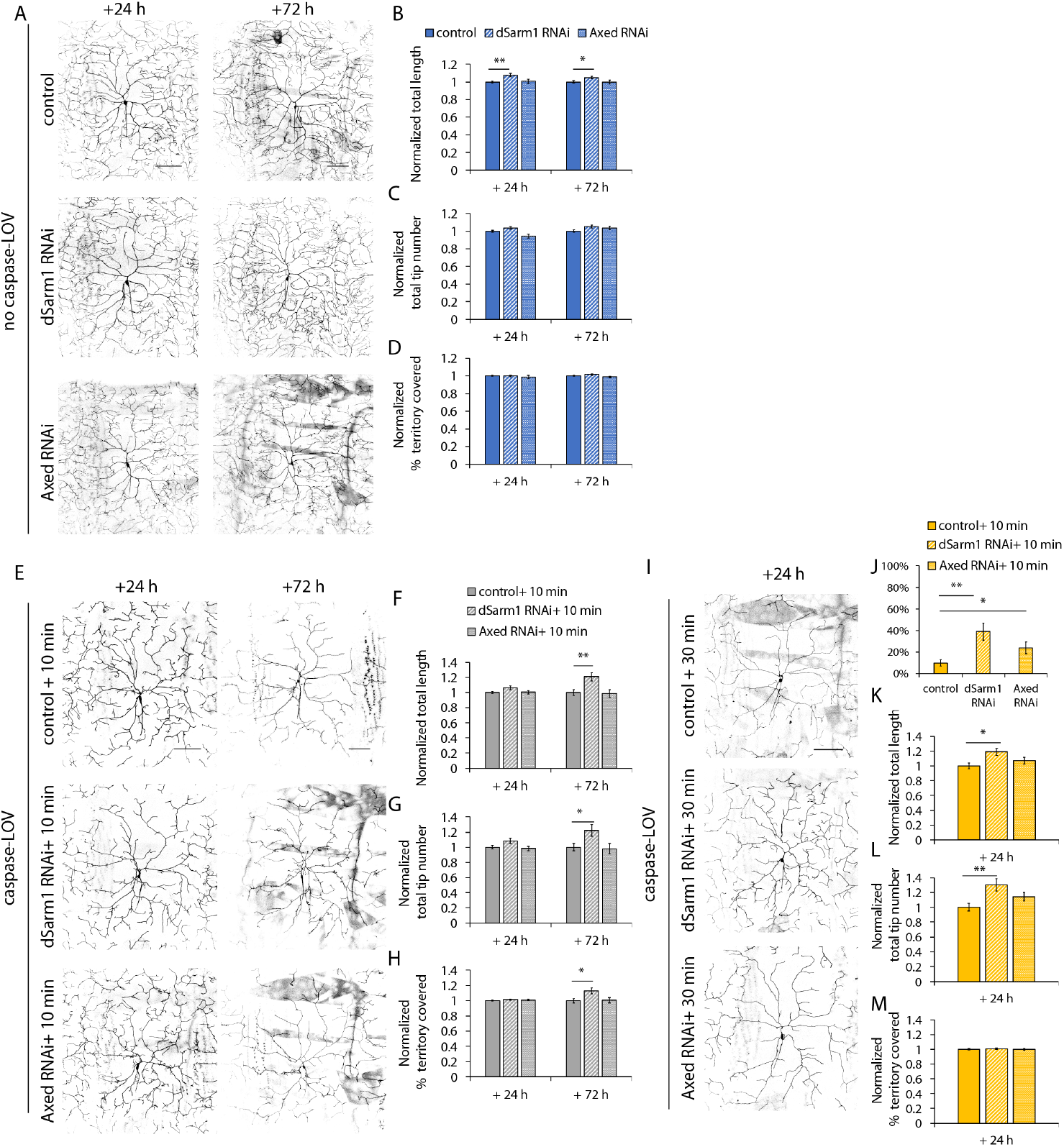
dSarm1 knockdown improves neuronal survival and allows neurons to retain longer dendrites throughout development and upon caspase-3 induced degeneration, while Axed knockdown only increases neuronal survival. (A) Representative images of c4da neurons labeled by ppk-tdGFP with ppk-Gal4 driving expression of UAS-luciferase (control), UAS-dSarm1 RNAi, or UAS-Axed RNAi. C4da neurons expressing dSarm1 RNAi have longer dendrites at early development (+ 24 h) and the dendrite length remains long until late development (+ 72 h). Knockdown of Axed in c4da neurons does not affect dendrite development. (B-D) Quantifications of dendrite structures, including normalized length (B), normalized tip numbers (C), and normalized percentage of territory covered (D) of c4da neurons. (E, I) Representative images of c4da neurons expressing ppk-tdGFP and UAS-caspase-LOV and UAS-luciferase (control), UAS-dSarm1 RNAi, or UAS-Axed RNAi driven by ppk-Gal4. Larva were kept in the dark and illuminated for 10 min (E) or 30 min (I) at 48 h after egg laying and imaged after 24 h or 72 h. (F-H) Quantifications of dendrite structures, including normalized length (F), normalized tip numbers (G), and normalized percentage of territory covered (H) of c4da neurons illuminated for 10 min. (J-M) Quantifications of survival rate (J) and dendrite structures, including normalized length (K), normalized tip numbers (L), and normalized percentage of territory covered (M), of c4da neurons illuminated for 30 min. Scale bars =100 μm. * p<0.05, ** p<0.01, *** p<0.001, one way ANOVA with Tukey’s post hoc test for multiple comparison in (B-D, F-H, K-M), Kruskal-Wallis rank sum test with Dunn’s post hoc test further adjusted by the Benjamini-Hochberg FDR method for multiple independent samples (J); Error bars represent ± SEM. n ≥ 29 neurons for each experimental condition and timepoint.

### Wld^S^ can partially rescue caspase 3-induced neurodegeneration and impairment of thermal nocifensive behavior

The chronic low-level caspase-LOV activity in the dark caused mild but significant dendrite degeneration during early larval development (***Figure 1A***,***B***,***D***,***E***,***F***,***G***,***H***). This mild degeneration continued throughout development up to the stage of wandering larvae (***Figure 7A***). These c4da neurons displayed impaired dendrite structure including shorter dendrites, fewer dendrite tips, and a lower percentage of territory covered (***Figure 7A***,***B***). We wondered whether these neurons with dendrite degeneration can fulfill normal sensory function. As nociceptive neurons, c4da neurons are required for the aversive rolling behavior when larvae encounter nocifensive stimuli such as high temperature (Babcock et al., 2009; Hwang et al., 2007). To test whether caspase-3 induced neurodegeneration affects the neuronal function, we examined the thermal nocifensive behavior in wandering larvae kept in the dark with or without caspase-LOV expression at two nocifensive temperatures, 46°C for tests of insensitivity, and 42°C for testing hypersensitivity (Honjo et al., 2016). We measured the time it took for an individual larva to initiate the rolling behavior within 20 seconds (s) of contacting the thermal probe at high temperature. We also quantified the percentage of non-responders (larvae that did not respond within 20 s). We found that larvae kept in the dark with chronic low-level caspase-LOV activity in c4da neurons took longer to initiate rolling behavior to escape the high temperature and a higher percentage of them were non-responders (***Figure 7C***,***D***).

**Figure 7.**
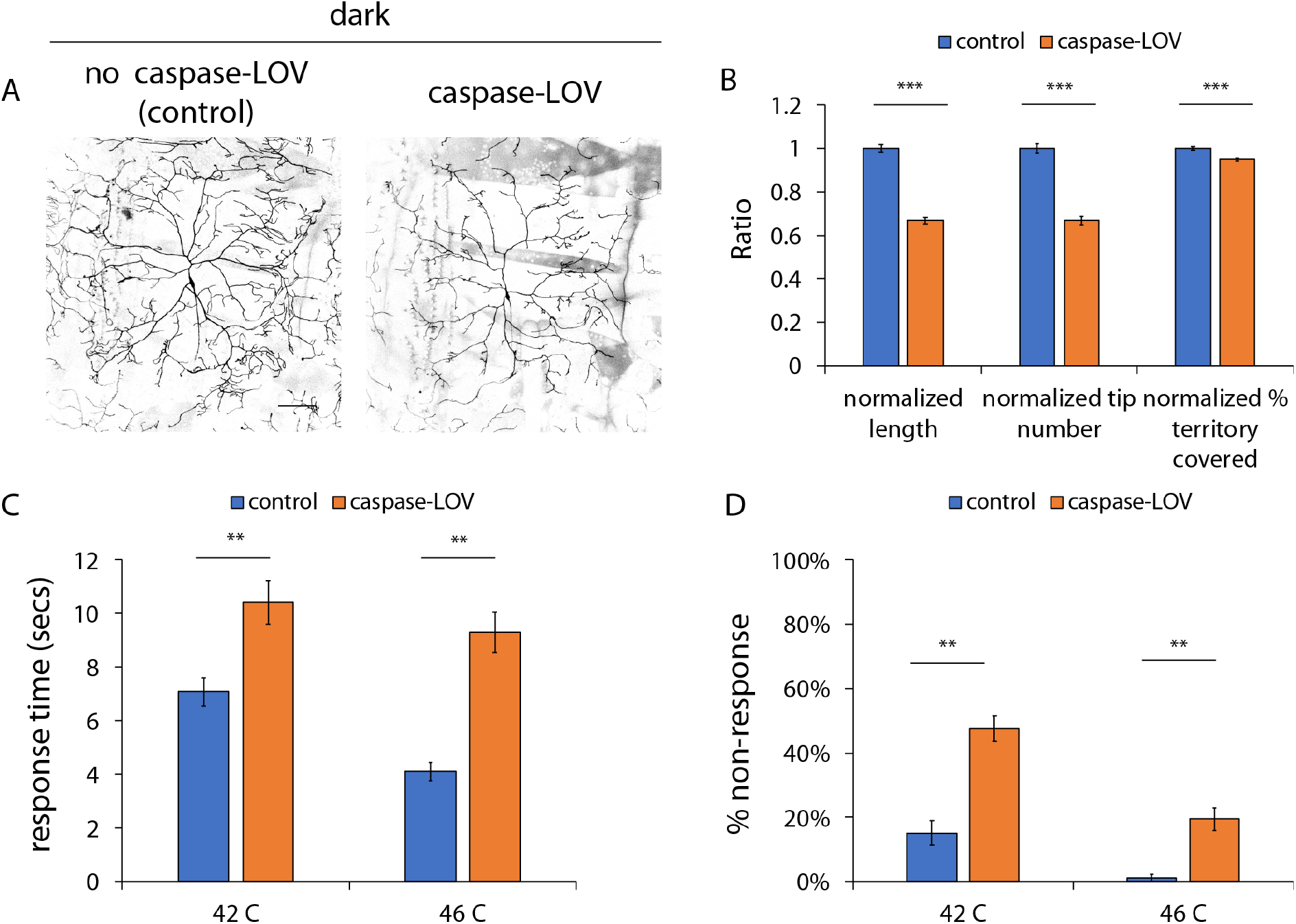
Activation of caspase-LOV in c4da neurons impaired the thermal nociceptive behavior. (A) Representative images of c4da neurons expressing UAS-tdTOM and UAS-luciferase (control) or UAS-tdTOM and UAS-caspase-LOV (caspase-LOV) driven by ppk-GAL4. Larvae are raised in the dark. C4da neurons expressing caspase-LOV have significant reductions in dendrite length, tip numbers and percentage of territory covered compared to control neurons at third-instar wandering stage. (B) Quantifications of dendrite structures, including normalized length (left), normalized tip numbers (middle), and normalized percentage of territory covered (right) of c4da neurons. (C-D) Aversive responses of third-instar wandering larvae in response to nocifensive temperature at 42°C and 46°C is affected by low-level caspase-LOV activation in the dark with longer response times (C) and a higher percentage of non-responders (D). Animals were classified as ‘‘non-responder’’ if the larva did not initiate the rolling behavior within 20 s of heated thermal probe touching the body wall. Scale bars =100 μm. * p<0.05, ** p<0.01, *** p<0.001, Student’s *t* test in (B-D). Error bars represent ± SEM. B: n = 31 (control) or 39 (caspase-LOV) neurons were tested. C-D: n ≥ 85 animals were tested for each genotype and temperature.

Having found that Wld^S^ expression in c4da neurons afforded protection from caspase-3 induced dendrite degeneration, we tested for its effect on the caspase-3 induced deficiency in the thermal nocifensive behavior. Without caspase-LOV, Wld^S^ expression in c4da neurons slightly increased the number of dendrite tips of c4da neurons in the wandering larvae (***Figure 8A***,***B***) but did not change their response time or the percentage of non-responders in the thermal nocifensive behavior (***Figure 8C***,***D***). We then examined the thermal nocifensive behavior of wandering larvae expressing caspase-LOV along with UAS-mIFP-2A-HO1 (control) or UAS-Wld^S^ (Wld^S^). With chronic low-level caspase-LOV activity in the dark, Wld^S^ expression resulted in longer and more numerous dendrite tips but with a smaller percentage of territory covered by dendrites (***Figure 8E***,***F***). Moreover, Wld^S^ partially rescue the caspase-3 induced impairment in thermal nocifensive behavior. Wld^s^ expression in c4da neurons reduced the averaged time to respond to a 46°C heat probe (***Figure 8G***). It also reduced the percentage of non-responders in larvae expressing caspase-LOV and kept in the dark (***Figure 8H***). These findings indicate that Wld^S^ expression in c4da neurons not only afforded preservation in dendrite structures but also protected neuronal functions critical for behavioral response to nociceptive stimuli.

**Figure 8.**
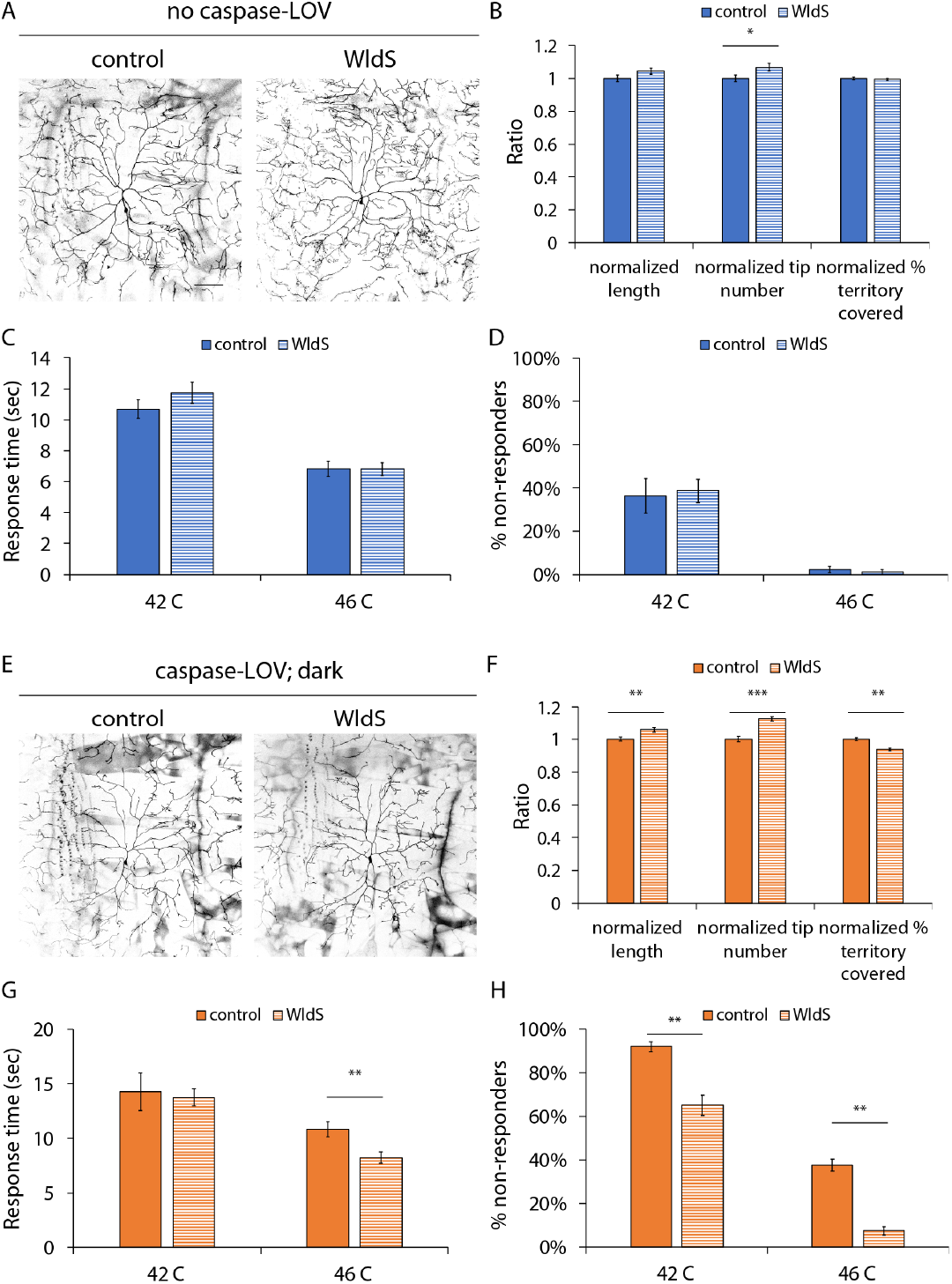
Wld^S^ can reduce caspase-3 induced dendrite degeneration and impairment in the thermal nocifensive behavior. (A) Representative images of c4da neurons expressing tdTOM and UAS-mIFP-2A-HO1 (control) or UAS-tdTOM and UAS-Wld^S^ (Wld^S^) driven by ppk-GAL4. Larvae are raised in the dark. C4da neurons expressing Wld^S^ have significant increases in total dendrite tip numbers compared to control neurons during dendrite development at the third-instar wandering stage. (B) Quantifications of dendrite structures, including normalized length (left), normalized tip numbers (middle), and normalized percentage of territory covered (right) of c4da neurons. (C-D) Wld^S^ expression on its own in c4da neurons does not change the response time (C) nor the percentage of animals that do not respond to nocifensive temperatures of 42°C and 46°C (D). (E) Representative images of c4da neurons expressing tdTOM, and caspase-LOV and UAS-mIFP-2A-HO1 (control) or UAS-tdTOM, UAS-caspase-LOV, and UAS-Wld^S^ driven by ppk-GAL4. Larvae are raised in the dark. C4da neurons expressing Wld^S^ can protect neurons from dendrite degeneration induced by low-level caspase-LOV activation in the dark as shown by significant increases in both dendrite length and tip numbers. There are reductions in the percentage of territory covered in Wld^S^ expressing c4da neurons compared to control neurons. (F) Quantifications of dendrite structures, including normalized length (left), normalized tip numbers (middle), and normalized percentage of territory covered (right) of c4da neurons. (G-H) Slower thermal nocifensive response induced by caspase-3 can be partially rescued by expression of Wld^S^ in c4da neurons. Wld^S^ expression in animals with low-level caspase-LOV activation in the dark leads to a decreased response time (G) at 46°C and the lower percentage of non-responding animals (H) in response to nocifensive temperature at 42°C and 46°C. Scale bars =100 μm. * p<0.05, ** p<0.01, *** p<0.001, Student’s *t* test in (B-D, F-H). Error bars represent ± SEM. B, F: n ≥ 51 neurons for each genotype. C-D, G-H: n ≥ 75 animals were tested for each genotype and temperature.

While knockdown of dSarm1 or Axed did not affect dendrite structure of c4da neurons in the wandering larvae (***Figure 9A***,***B***), dSarm1 knockdown delayed the behavioral response to contacts with a probe heated to 46°C (***Figure 9C***). Knockdown of either dSarm1 or Axed increased the population of non-responders upon encounter with a probe at the nocifensive temperature of 42°C (***Figure 9D***). With mild degeneration induced by the chronic low-level caspase-LOV activity in the dark throughout larval development, dSarm1 knockdown caused a small increase in the percentage of territory covered by c4da neuron dendrites in the wandering larvae (***Figure 9E***,***F***), while RNAi knockdown of Axed did not affect the degeneration of dendrite structure (***Figure 9E***,***F***). Notably, knockdown of dSarm1 or Axed reduced the thermal nocifensive behavior of larvae with caspase-3 induced neurodegeneration (***Figure 9G***,***H***). Larvae with dSarm1 knockdown in c4da neurons took longer to avoid the probe heated to 42°C (***Figure 9G***). Knockdown of dSarm1 or Axed in c4da neurons increased the percentage of non-responders when stimulated with a probe heated to 42°C or 46°C (***Figure 9H***). These results indicate that knockdown of dSarm1 or Axed in c4da neurons impaired the thermal nocifensive behavior of larvae during development and further exasperated the deficient thermal nocifensive behavior owing to caspase-3 induced degeneration of c4da neurons.

**Figure 9.**
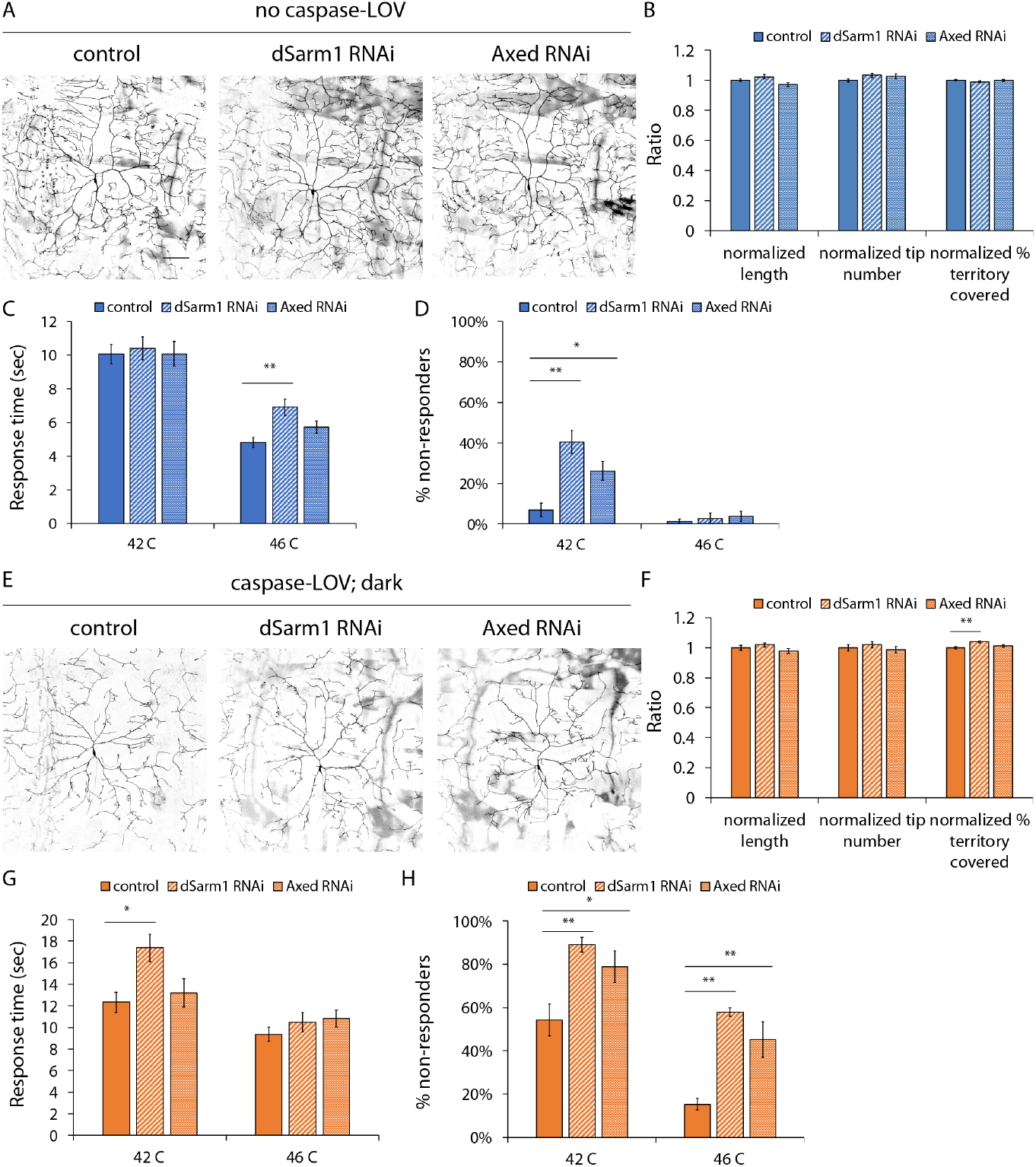
Knockdown of dSarm1 or Axed reduces the thermal nocifensive behavior. (A) Representative images of c4da neurons expressing UAS-tdTOM and UAS-luciferase (control), UAS-tdTOM and UAS-dSarm1 RNAi (dSarm1 RNAi), or UAS-tdTOM and UAS-Axed RNAi (Axed RNAi) driven by ppk-GAL4. Knockdown of dSarm1 or Axed by RNAi in c4da neurons does not change dendrite structures compared to control neurons during dendrite development at the third-instar wandering stage. All larvae are raised in the dark. (B) Quantifications of dendrite structures, including normalized length (left), normalized tip numbers (middle), and normalized percentage of territory covered (right) of c4da neurons. (C-D) Third-instar wandering larvae expressing dSarm1 RNAi in c4da neurons responded slower (C) and had a higher percentage of non-responding animals at 42°C (D). Knockdown of Axed also increases the percentage of animals with no response to 42°C(D). (E) Knockdown of dSarm1 or Axed by RNAi in c4da neurons does not change dendrite degeneration induced by caspase-LOV activation in the dark compared to control neurons at the third-instar wandering stage. (F) Quantifications of dendrite structures, including normalized length (left), normalized tip numbers (middle), and normalized percentage of territory covered (right) of c4da neurons. (G-H) Slower thermal nocifensive response induced by caspase-3 is worsen when dSarm1 and Axed are knocked down in c4da neurons. When dSarm1 expression is knocked down in these neurons, there is an increased response time at 42°C (G). When either dSarm1 or Axed are knocked down, there is a higher percentage of non-responding animals when probed at 42°C and at 46°C (H). Scale bars =100 μm. * p<0.05, ** p<0.01, *** p<0.001, one-way ANOVA with Tukey’s post hoc test for multiple comparison in (B-D, F-H). Error bars represent ± SEM. B, F: n ≥ 44 neurons for each genotype. C-D, G-H: n ≥ 72 animals were tested for each genotype and temperature.

## DISCUSSION

In this study, we established a new neurodegeneration and repair assay system with the photo-switchable caspase-3, caspase-LOV, to elucidate the mechanisms underlying dendrite degeneration and repair. To characterize the caspase-3 induced neurodegeneration, we focused on the dendrite morphology for different classes of da neurons and observed cell type-specific cellular responses. We also examined the c4da neurons-mediated thermal nocifensive behavior to reveal the functional consequence of neurodegeneration. We found that Wld^S^, a key molecule involved in the Wallerian axon degeneration, can protect dendrite structure and reduce the impairment of thermal nocifensive behavior caused by caspase-LOV activation in c4da neurons. Knockdown of dSarm1 reduced the caspase-3 induced loss in dendrite structure, whereas knockdown of Axed did not affect dendrite degeneration. Knockdown of dSarm1 or Axed led to impaired thermal nocifensive behavior with or without caspase-3 induced degeneration of c4da neurons. Along with the previously established laser severing injury model, our new model with adjustable caspase-LOV activation provides a useful platform to identify regulators and to improve our understanding of dendrite degeneration and repair.

### Cell type-specific cellular responses upon caspase-3 induced dendrite degeneration

We examined how three different classes of da neurons react to caspase-LOV activation and found cell type-specific responses to caspase-3 induced dendrite degeneration. Dendrites of c1da, c3da and c4da neurons all exhibit more severe degeneration following longer duration of caspase-LOV activation. The survival rates of these three classes of da neurons also decreased with longer caspase-LOV activation. However, these da neurons differ in the dynamic dendrite changes over the 24-72 h period following illumination. Remarkably, c1da neurons displayed an increased percentage of added dendrite tips number following 30 min but not 2 h of caspase-LOV activation compared to neurons kept in the dark. This reactivation of the growth program is exhibited by c1da neurons but not c3da or c4da neurons. The c3da neurons continued to grow in dendrite length and tip numbers over the 24-72 h period following 30 min and 2 h of caspase-LOV activation, as control c3da neurons did. In contrast, caspase-LOV activation for 2 h induced significant changes in dendrite length and tip numbers in both c1da and c4da neurons. Unique to c4da neurons is an increase in the percentage of added dendrite tips with caspase-LOV activity in the dark.

Extensive studies in da neurons revealed the cell-type specific dendrite morphology, gene expressions, dendrite remodeling and injury responses (Grueber et al., 2002; Jan and Jan, 2010; Shimono et al., 2009; Song et al., 2012; Thompson-Peer et al., 2016). Our data further suggest that different classes of da neurons are also equipped with specialized mechanism to handle caspase-3 induced neurodegeneration. By including different classes of da neurons in our study, we aim for a more comprehensive survey of how to protect dendrites from degeneration and improve recovery of neuronal functions. For example, future studies of c1da neurons could elucidate the growth programs reactivated following degeneration and assess whether such programs can be transferred to other cell types. As to c3da neurons, it would be of interest to investigate how they can withstand the caspase-LOV activation without halting their growth.

### Protection afforded by Wld^S^ may vary with the degree of neurodegeneration

The Wallerian degeneration pathway important for axon degeneration serves as a prominent target for therapy. In this study, we focused on the impacts of neurodegeneration on dendrites, which together with axons are responsible for maintaining neuronal functions. Our study of the impact of caspase-3 induced neurodegeneration on dendrite morphology and thermal nocifensive behavior reveals intriguing involvement of the Wallerian degeneration pathway. We found that with 10-30 min caspase-LOV activation, Wld^S^ can partially rescue the caspase-3 induced deficiency in dendrite structure and neuronal survival. Wld^S^ also afforded protection for the impaired dendrite structure and thermal nocifensive behavior caused by the chronic low-level caspase-LOV activity in the dark. Interestingly, with continuous activation of the photo-switchable caspase-3 via illumination for days, a much stronger perturbation employed in a previous study, Wld^S^ fails to rescue the survival of flies with neuronal expression of caspase-LOV (Smart et al., 2017). This suggests that the ability of Wld^S^ to provide protection may depend on the level of caspase-LOV activation in a neuron. Whether different mechanisms are used for dendrite degeneration or repair in neurons experiencing different levels of caspase-LOV activation is an interesting question that can be explored using this tunable neurodegeneration model in the future.

### Multiple roles of Wld^S^ in caspase-3 induced degeneration and repair

The preservation of neuronal function by Wld^S^ following caspase-LOV activation may result from the retained dendrite structures and/or axons, or a complex combination of different factors. In this study, we did not examine the caspase-3 induced axon degeneration and repair. The axons of the da neurons project deep into the ventral nerve cord and connect with central neurons to form circuits required for the avoidance behavior. These axons form bundles, while the dendrite arbors of da neurons display readily discernible patterns. To study caspase-3 induced axon degeneration and repair, future studies could examine cell types more suitable for imaging the axon morphology with established axon-dependent functional readouts, such as wing neurons or olfactory receptor neurons (ORNs) (Neukomm et al., 2017; Osterloh et al., 2012).

### dSarm1 and Axed play different roles in dendrite development, caspase-3 induced dendrite degeneration, and the thermal nocifensive behavior

With recent advances in the understanding of the Wallerian degeneration pathway, additional regulators have been identified, including dSarm1 and Axed. Studies in *Drosophila* and mice suggest that dSarm1 acts downstream of Wld^S^ while Axed may be either downstream of dSarm1 or involved in a separate pathway (Coleman and Höke, 2020; Neukomm et al., 2017; Osterloh et al., 2012; Sambashivan and Freeman, 2021). In this study, we examined the roles of dSarm1 and Axed in dendrites and found that their functions diverged from those of Wld^S^ during dendrite development, caspase-3 induced dendrite degeneration, and the thermal nocifensive behavior.

Previous studies report that knockout of dSarm1 specifically in c4da neurons does not affect dendrite structure but can protect c4da neurons from injury and PS-induced dendrite degeneration in wandering larvae during late larval development (Ji et al., 2021). Knockout of Axed in c4da neurons partially affects dendrite degeneration induced by injury but does not alter the degeneration in response to PS exposure (Ji et al., 2021). In this study, we found that knockdown of dSarm1 but not Axed in c4da neurons led to longer dendrites during early dendrite development and following caspase-3 induced dendrite degeneration. However, reduced dSarm1 expression in c4da neurons did not protect them against caspase-3 induced impairments in their dendrite structure and neuronal functions later in the development during the wandering stage. At the behavioral level, we found that knockdown of dSarm1 or Axed in c4da neurons of control larvae with or without caspase-LOV impaired the thermal nocifensive behavior. Thus, dSarm1 and Axed affect c4da neurons-mediated thermal nocifensive behavior of larvae without altering dendrites of c4da neurons.

### Functions of dSarm1 and Axed in other *Drosophila* neurons and in mammalian neurons

Apart from c4da neurons, roles of dSarm1 and Axed in dendrite morphology and neuronal functions in other cell types have been described. Sarm1 knockdown in cultured hippocampal neurons or in mice results in simplified dendrite structure instead of longer dendrites as we observed in c4da neurons (Chen et al., 2011). In the mushroom body gamma neurons of the fly central nervous system, dSarm1 and Axed mutations do not affect dendrite pruning (Neukomm et al., 2017; Osterloh et al., 2012). For the behavioral functions, Sarm1 knockdown in mice causes deficiency in associative memory, cognitive flexibility and social interactions (Lin and Hsueh, 2014), whereas flies containing dSarm1 or Axed mutant Johnston’s organ (JO) clones can still elicit JO neurons-mediated grooming behavior (Neukomm et al., 2017). It thus appears that the functions of these proteins may vary with their subcellular localization, the cell types, the time in development, as well as the species.

### Advantages of the new model for caspase-3 induced neurodegeneration

The range of dendrite degeneration and repair resulting from varying degrees of caspase-LOV activation demonstrates the versatility of the photo-switchable caspase-3 system to induce degeneration in *Drosophila* da neurons. This new model has several strengths. First, with photo-switchable caspase-3, the timing and degree of degeneration can be controlled by adjusting the length and intensity of illumination. The activation of caspase-LOV lasts for the duration of the illumination and is reversible. In conjunction with the genetic tools available, we could induce degeneration in specific cell types. Finer spatial control may be achieved by locally illuminating certain areas of the cell viewed under the microscope or by targeting the photo-switchable caspase-3 with linked peptide sequences to specific subcellular compartments. Second, in contrast to laser severing of dendrites, the photo-switchable caspase-3 allows for infliction of neuronal injury systematically in a way that is considerably less labor intensive. It is thus amenable to screens of genetic manipulations or pharmacological drug libraries to dissect the underlying cellular and molecular mechanisms. Third, this model is physiologically relevant, given that caspase-3 plays a role in the developmental pruning of axon and dendrite (Kuo et al., 2006; Williams et al., 2006; Schoenmann et al., 2010) as well as axon degeneration initiated by trophic factor withdrawal (Nikolaev et al., 2009; Schoenmann et al., 2010, Simon 2012). The discoveries made possible with the photo-switchable caspase-3 system will therefore be likely to yield information about physiologically relevant neurodegeneration that occurs during developmental pruning, trophic factor withdrawal, and disease. Our model can complement the existing injury models, including laser ablation-induced dendrite degeneration and PS-induced dendrite degeneration (Sapar et al., 2018; Tao and Rolls, 2011) and provide an alternative route to study how to repair dendrites following neurodegeneration. It is currently unclear whether neurons respond to different insults the same way or whether insult-specific response pathways exist. In order to develop effective therapies, it is important to investigate how neurons respond to different types of injuries.

### Possible improvements of the new model

In this study, we set up a degeneration and repair model for larval sensory neurons. The repair process identified in the larval sensory neurons could be a combination of developmental growth and a repair response specific to caspase-3 induced degeneration. To focus on the contribution from the repair process and to identify ways to re-establish the growth capacity of neurons, it is desirable to extend the system to the adult sensory neurons. The adult sensory neurons reach maturity around 3 days after eclosion and have stabilized dendrite structure throughout adulthood (DeVault et al., 2018). Therefore, dendrite elongation or addition following caspase-3 induced degeneration in adult fly would correspond to regeneration and repair.

In this proof-of-principle initial study, transgenes and RNAis are expressed before caspase-LOV activation, so the effects could be due to prevention of damage or repair of damage. Future improvements for better temporal control could make use of either a pharmacologically controlled gene switch system or the temperature-sensitive GAL80 repressor (Gal80ts) (Nicholson et al., 2008; Zeidler et al., 2004). Whereas we focused on cell autonomous factors in this study, we recognized there are likely non-cell autonomous contributions from epidermal cells and glial cells (DeVault et al., 2018; Liu and Jan, 2020; Song et al., 2012; Yadav et al., 2019; Yin et al., 2021). Future studies of dendrite degeneration at different stages of development as well as adulthood may shed light on strategies to prevent neurodegeneration, to diagnose neurodegeneration early, and to develop drugs promoting neural recovery from injury and diseases.

## METHODS

### Fly stocks and genetics

Animals were reared at 25°C or at 22°C for monitoring the dendrite degeneration and repair. The fly strains used in this study were as follows: UAS-Wld^S^ (a generous gift from Ashley Smart at UCSF (Hoopfer et al., 2006)), Gal4^19-12^ (Xiang et al., 2010), Gal4^2-21^ (Grueber et al., 2003a), ppk-Gal4 (Grueber et al., 2003b), ppk-CD4-tdGFP (Han et al., 2011), UAS-caspase-LOV (BL76355, a generous gift from Ashley Smart at UCSF), UAS-tdTomato (Han et al., 2011), UAS-mIFP-T2A-HO1 (attp40 on 2^nd^ chromosome used in this study. a generous gift from Xiaokun Shu, UCSF), UAS-luciferase (BL35788, control RNAi for the TRiP lines) UAS-dSarm1-RNAi (BL 63525), UAS-Axed-RNAi (BL 62989). The RNAi lines we used in the study are all VALIUM20-series TRiP RNAi fly stocks that produce short hairpin RNAs (shRNAs) and give stronger knockdown efficiency then VALIUM10-series TRiP RNAi flies (Ni et al., 2011). The tester lines for RNA interference (RNAi) or overexpression experiments was ppk-gal4, ppk-CD4-tdGFP; UAS-caspase-LOV. RNAi or overexpression experiments were performed by crossing the tester lines to the variety of transgenic fly strains. To control for caspase-LOV expression dosage in different genotypes, we used UAS-mIFP-T2A-HO1 (wild-type, w^1118^) as control for Wld^S^ experiments and UAS-luciferase (yv flies) as control for RNAi experiments. We found slight differences for thermal nocifensive behavior in genotypes, so we used different fly strains as controls for Wld^S^ and RNAi lines.

### Illumination box with LED strips

We collect eggs laid in the dark for 2 h and kept them in the dark at 25°C until illuminated at 48 h after egg laying (AEL). To activate the photo-switchable caspase-3, freely moving larvae were picked and transferred the transparent agar plates with a thin layer of yeast. Larvae were moved back to yeasted grape plate and kept in the dark at 22°C after different durations of blue LED illumination. Lower raising temperature to 22°C can delay development and increase the temporal resolution of the repair process following caspase-3 activation. To avoid lights, grape juice plates are store in 10 mm petri dishes wrapped with foil. A homemade 40 cm x 10 cm x 15 cm carbon box was used to shield larvae from ambient light and to house three 10cm long and 8mm wide Blue 3528 LED strip, (peak at 460nm, Environmental Lights) stick on the ceiling of the box in parallel and connected by wires. LED strips are wired to a connector with DC jack (Environmental Lights) and then a LED Power Supply Adapter (HitLights). The power of the light 15 cm away from the LED strips, where larvae were kept, is 0.91 mW/cm^2^.

### *In vivo* time lapse imaging

Live imaging was performed as described (Emoto et al., 2006; Parrish et al., 2007). Larvae were anesthetized with diethyl-ether for 5-8 minutes (Acros Organics) before mounted in glycerol on top of a thin patch of agarose. After images were acquired using a Leica SP5 microscope with a 20X oil objective (NA 0.75), larva was returned to yeasted grape juice agar plates or sacrificed if this is the end of imaging timepoints. Sum slices for Z-projection were generated using ImageJ software and used for dendrite structure prediction as described later.

To visualize neurons, c4da ddaC neurons were labeled by expressing UAS-CD4-tdTOM using the ppk-GAL4 driver or by using the direct fusion line ppk-tdGFP. C1da ddaE neurons are visualized through mCD4-tdTOM driven by Gal4^2-21^. C3da ddaF neuron with UAS-CD4-tdTOM driven by the GAL4^19-21^ along with Repo-Gal80 to eliminate the expression in glial cells (Awasaki et al., 2008; Xiang et al., 2010)

### Deep learning based-automatic dendrite structure prediction

We utilized two methods to segment the dendrite structures of the da neurons for morphological quantification. For ddaE, c1da neurons, and ddac, c4da neurons, in Figure 1, we reconstructed individual neurons using Vaa3D-Neuron 2.0: 3D neuron paint and tracing function in Vaa3D (http://vaa3d.org/) with manual correction and validation of the tracing (Peng et al., 2010).

For the rest of ddaC neurons in this study, we established a U-Net based deep learning model for automatic dendrite structure segmentation which produces segmentation maps with pixel intensity representing the probability of dendrite structure. We followed the U-Net architecture specified in the original study (Ronneberger et al., 2015) with modifying the channel number of the final segmentation map from 2 to 1 since we only predicted dendrite structure versus background. Each training data consisted of a maximum intensity Z-projection image of one neuron manually cropped by drawing a ROI, paired with the manually segmented dendrite structure (mask) generated using the plugin, “simple neurite tracer”, in ImageJ. In total, we generated 29 sets of image-mask pairs for training and 8 sets for validation with datasets generated in-house. Two data augmentation strategies were used to increase the model robustness. First, an area of 512×512 pixels was randomly cropped from each input 1024×1024 training image and the associated mask. Then the cropped image and mask were randomly flipped horizontally and vertically with probability 0.5. We used the sum of binary cross-entropy and Dice loss (defined as 1 – Dice coefficient) as the loss function and trained the model with Adam optimizer at learning rate 1e-4 for 500 epochs. The best model evaluated by Dice loss using the validation dataset was chosen for the downstream analysis. Our best model achieved the Dice loss at 0.13 and 0.16 for training and validation datasets, respectively.

A threshold of 0.5 was used to binarize segmentation maps generated by the model. We found high correlation (R^2^ = 0.98) in total dendrite length of larval neurons between model-predicted segmentation and manual reconstruction, while tip numbers only showed moderate correlation (R^2^ = 0.45). This was because tip number was more sensitive to the discontinuity and small fragments occasionally found in model-predicted segmentation masks. Therefore, we included a 3-step post-processing procedure to exclude small fragments and reduce the discontinuity in the segmented dendrite structure. First, small objects with area less than 10 pixels were discarded. Second, dilation with a cross-shaped structuring element (connectivity=1) was used to fill in the gaps. Finally, skeletonization using the *skeletonize* function from Python scikit-image package was applied to obtain the final segmentation for the downstream morphology quantification. With post-processing to fill in gaps and remove small fragments, we observed a dramatic increase in the correlation of tip numbers (R^2^ = 0.97) and a slight increase for total dendrite length (R^2^ = 0.99).

This system can be applied to predict the structures of other type of neurons either using the exiting models or retain models with new set of training datasets. One limitation is to separate the individual neurons at the manual ROI selection step. For example, the Gal4^19-12^ and Gal4^2-21^ drivers sometimes have weak expression in surrounding neurons which is hard to separate. When the neurons are also well-marked by fluorescence proteins, they can be recognized by the prediction model and included as part of the c1da neurons which introduce false positive errors. Therefore, we did not use the model for the c1da neurons.

### Quantification of dendrite structure

With the prediction model described above along with the post-processing python code, we can obtain the total dendrite length, total dendrite tip numbers and skeletal images of predicted dendrite structures. Using the skeletal images, we performed Sholl analysis of dendrite branches to determine the complexity of the dendrite structure. The crossing continuous circles were separated by 0.76μm on either manually traced or predicted dendrite arbors. To determine percentage of territory covered, we measured the area of dendrite arbor of neuron of interest covered and divided it to total area of the hemisegment of the body wall. The territory covered is measured using ROI selection tools in ImageJ. We defined a cell as “survived” if the average dendrite length (total dendrite length/total tip numbers) over 10 μm for c4da neurons. For c3da and c1da neurons, we identified neurons with more than 2 dendrite tips (more than one dendrite branch) as survived. To reduce the batch-by-batch variations, we normalized the quantifications to the controls for each batch before combining all data. For comparison between different conditions, the number was normalized to the averaged number in dark (control). The results are normalized to the controls for each set of experiments before combining.

### Thermal nocifensive behavior

For thermo-nociception using a local hot probe, a custom-built thermo-couple device was used to keep the applied temperature constantly at 42 or 46 °C as desired. Stage and density-controlled 3rd instar wandering stage larvae were used. Freely moving larvae were touched with the hot probe on mid-abdominal segments until the execution of nociceptive rolling avoidance behavior. Animals were monitored under cell phone camera (Nokia 6.1) and the time it takes to initiate the rolling behavior for high temperature were counted with in 20 s. The animals that take longer than 20 s to response were classified as no responders. The percentages of no responder were calculated by dividing numbers of no responders by numbers of total tested animals. Each genotype was tested multiple times on different days and data from all trials was combined.

### Software

The code used for deep learning based automatic dendrite structure prediction is written in python/TensorFlow. We trained our model on a Quadro P5000 GPU with 16 GB RAM in a Dell Precision 7920 Tower with Dual Intel Xeon Gold 6136 CPUs (3.0/3.7GHz), having 12 cores and 128 GB RAM. The operating system was Windows 10. We have tested our system on Mac and Windows operating system. The software package, training and example testing images are available on the GitHub repository (https://github.com/chienhsiang/dendrite_U-Net).

### Statistical tests

All data are presented as mean ± standard error of the mean (SEM) based on at least three independent experiments. Data are considered significantly different when p values are less than 0.05. Student’s t test was used for comparisons of two groups. One-way ANOVA with Tukey’s post hoc test was used for comparisons of multiple groups. The Kruskal-Wallis rank sum test with Dunn’s post hoc test further adjusted by the Benjamini-Hochberg FDR method was used for multiple comparisons of nonparametric samples. Statistics analysis was performed and prepared using JASP (Version 0.14). All samples were prepared and analyzed in parallel.

## ACKNOWLEDGMENTS

We would like to thank members of the Jan Lab for helpful discussions. We want to thank Jacob Jaszczak, Caitlin O’Brien, Liying Li, and Ashely Smart for critical reading and suggestions on the manuscript. We are grateful for Ashley Smart and Xiaokun Shu at UCSF for kindly sharing fly stocks with us. We are grateful for Caitlin O’Brien for providing the training datasets. Research reported in this publication was supported by the National Institute of Neurological Disorders and Stroke (R35NS097227 to YNJ). Yuh-Nung Jan and Lily Y. Jan are investigators at the Howard Hughes Medical Institute.

## COMPETING INTERESTS

The authors declare no competing interests.

---

**Figure 1 – figure supplement 1.**
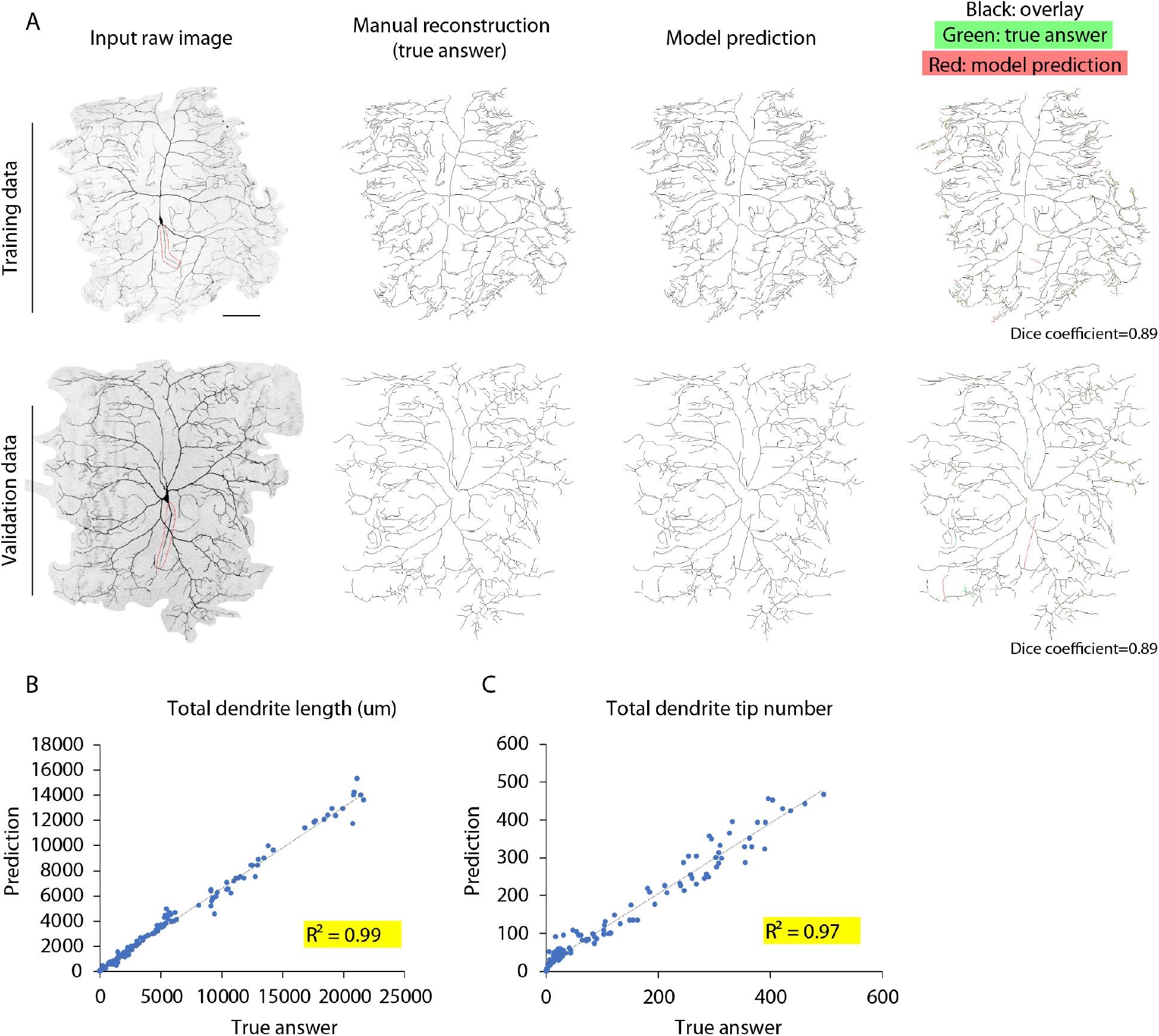
Deep learning-based automatic dendrite structure prediction. (A) Our in-house trained deep learning-based model performed well in dendrite segmentation. In the top row are images of a representative neuron from the training dataset and the bottom row is a neuron from the validation dataset (novel neurons for the model). The first column contains input Z-projection image of neurons manually cropped by drawing a ROI. Images in the second column are manually segmented dendrite structure (true answer) from ImageJ plugin, “simple neurite tracer”. Our model predictions are in the third column. The last column has overlay images from true answer and model prediction. The model reliably recognized most of the arbors as human as most of the dendrites are matched (marked in black) with few distal dim dendrites omitted by the model and only shown in the true answer (green) or only recognized by the model (red). Our model did not differentiate between axons and dendrites and sometimes counts the axon (circled with red dash line in the first column) as one of the dendrites (7 out of 37 neurons in the training and validation dataset). (B-C) Relationships between manual reconstruction (true answer) and the deep learning model (prediction) for total dendrite length and total dendrite tip number. After post-processing, our prediction model achieved 0.99 for R^2^ of total dendrite length (B) and 0.97 for R^2^ of tip numbers (C). Scale bars =100 μm. n = 160 neurons.

**Figure 5 – figure supplement 1.**
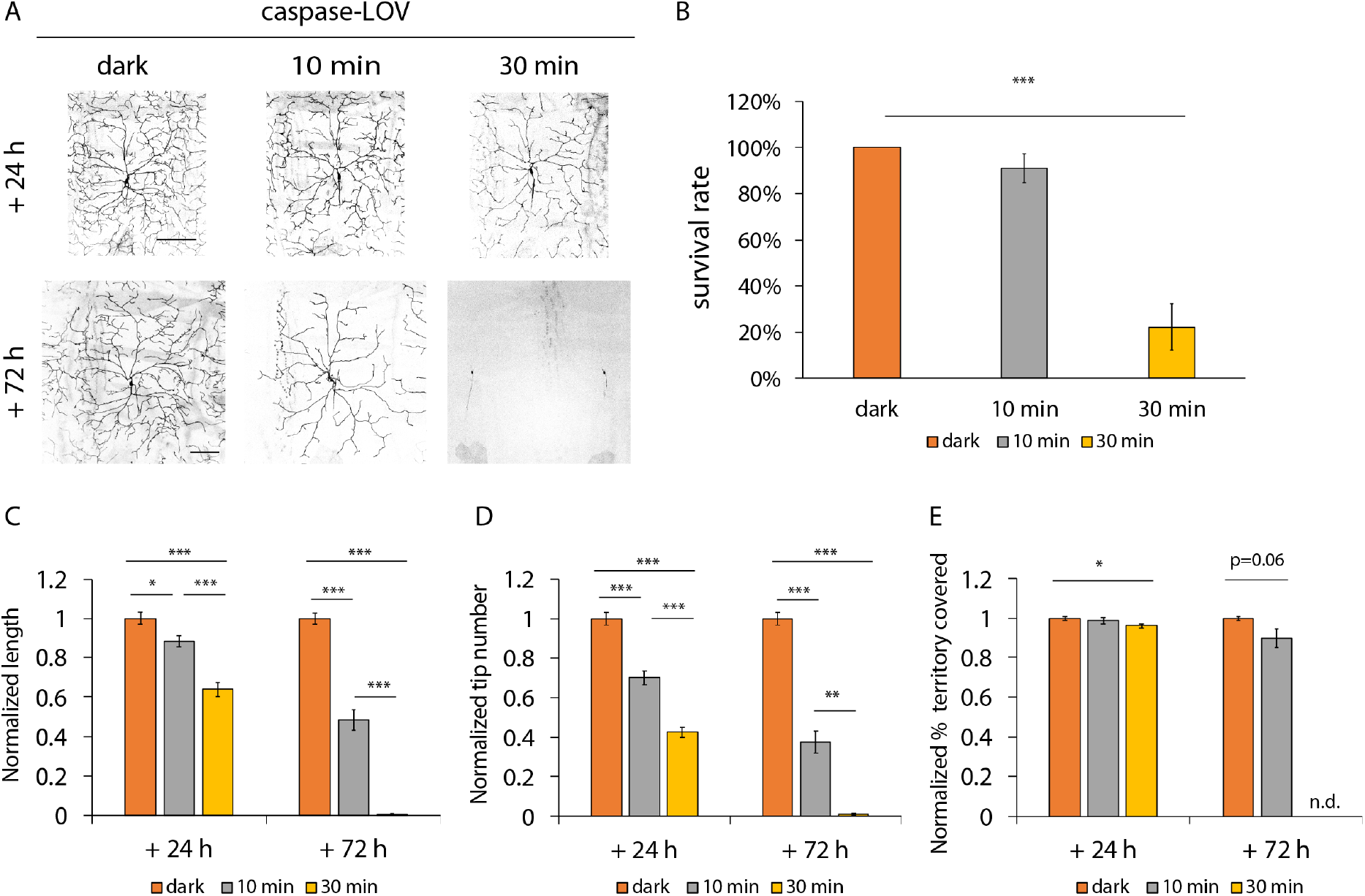
Degeneration and repair in the c4da neurons of tester animals. (A) Representative images of c4da neurons expressing UAS-luciferase and UAS-caspase-LOV driven by ppk-gal4 and labeled with ppk-tdGFP. Larvae were illuminated for 10 min or 30 min and imaged following the protocol in Fig. 1A. (B) Survival rates of c4da neurons decreased significantly upon 30 min illumination. (C-E) Quantifications of dendrite structures, including normalized length (C), normalized tip numbers (D), and normalized percentage of territory covered (E) of c4da neurons kept in the dark, illuminated for 10 min or illuminated for 30 min. The dendrite degeneration in the surviving c4da neurons is worse when illumination is extended. Scale bars =100 μm. * p<0.05, ** p<0.01, *** p<0.001, Kruskal-Wallis rank sum test with Dunn’s post hoc test further adjusted by the Benjamini-Hochberg FDR method for multiple independent samples (B); one-way ANOVA with Tukey’s post hoc test for multiple comparisons in (C-E). Error bars represent ± SEM. n = 16-24 neurons for each experimental condition and timepoint.

## Notes

### Competing Interest Statement

The authors have declared no competing interest.

